# HMCES DNA-protein cross-links promote template slippage during DNA replication

**DOI:** 10.64898/2026.08.14.744967

**Authors:** Xu He, Yixin Clem Xu, Yuge Chai, Kha T. Nguyen, Gaoyuan Liu, William A. Goddard, Daniel R. Semlow

## Abstract

During replication, nucleolytic processing of apurinic/apyrimidinic (AP) sites in single-stranded (ss)DNA is attenuated by the evolutionarily conserved 5-hydroxymethylcytosine binding, embryonic-specific (HMCES) protein. HMCES forms a covalent thiazolidine linkage with the ring-opened aldehyde form of a ssDNA AP site to stabilize the AP site and suppress the formation of DNA double-strand breaks. The resulting HMCES DNA-protein cross-link (DPC) can then be digested by the SPRTN protease and bypassed through mutagenic translesion synthesis (TLS). Here, we use *Xenopus* egg extracts and molecular dynamics simulations to investigate how HMCES-DPC formation influences the mutagenicity of AP site bypass. We show that SPRTN processes the HMCES-DPC to a five amino acid peptide adduct prior to TLS. Surprisingly, the mutagenicity of HMCES-DPC bypass is insensitive to the extent of DPC proteolysis and depends only on cross-link formation, which traps the AP site in a more dynamic ring-opened configuration. We further show that the spectrum of mutations produced during bypass of HMCES-adducts strongly depends on the template strand nucleotide immediately 5’ of the AP site. Our data support a model in which HMCES-DPC formation increases the conformational flexibility of the DNA template, allowing template slippage and use of the 5’ template nucleotide to direct insertion opposite the adducted AP site.

## Main

DNA apurinic/apyrimidinic (AP) sites, also known as abasic sites, are among the most abundant genomic lesions in cells. Recent studies suggest that a human cell may harbor hundreds to thousands of AP sites at steady state^1^. These lesions can arise through spontaneous depurination/depyrimidination or through the action of glycosylases that excise damaged or non-canonical nucleobases^2^. AP sites exist as an equilibrium mixture of ring-closed hemiacetals and ring-opened aldehydes^3^. The electrophilic aldehyde form of an AP site is highly labile and can react with primary amines to form cross-links with small-molecule metabolites, proteins and nucleic acids^4^. AP sites are also susceptible to β-elimination that results in single-strand DNA nicks or, in the case of clustered AP sites, highly toxic DNA double-strand breaks (DSBs).

AP sites are normally repaired by the base excision repair (BER) pathway^2^. An AP endonuclease incises the DNA backbone immediately 5’ of the lesion, generating a free 3’ hydroxyl terminus and 5’ deoxyribose phosphate (5’ dRP). Next, the 5’ dRP lyase activity of Pol β removes the 5’ dRP residue through a β-elimination-like mechanism, leaving behind a single nucleotide gap with a ligatable 5’ phosphate. Pol β then fills in the gap through templated DNA synthesis and DNA ligase seals the remaining nick to complete repair. Although BER of AP sites in double-stranded (ds)DNA is critical for genome maintenance, processing of AP sites in single-stranded (ss)DNA can instead promote genome instability. During replication, helicase-mediated unwinding of the DNA template can expose an AP site in ssDNA. Because the AP site lacks an instructional base, it also blocks nascent strand extension by replicative DNA polymerases, delaying the restoration of duplex DNA. Incision of an AP site in this ssDNA context by AP nucleases or AP lyases collapses the replication fork into a one-ended DSBs that can give rise to chromosomal aberrations. Because replication fork collapse is thought to be a major source of endogenous DSBs in mammalian cells^5^, proper regulation of AP site processing during replication is likely to be critical for preserving genome stability.

ssDNA AP sites are protected during replication by the 5-hydroxymethylcytosine binding, embryonic stem cell-specific (HMCES) protein^6–9^. The SOS response associated peptidase (SRAP) domain of HMCES, which is conserved across all domains of life, binds specifically to ssDNA and reacts with the ring-open aldehyde form of an AP site through an N-terminal cysteine to form a thiazolidine linkage (Figure 1A). The resulting HMCES DNA-protein cross-link (DPC) renders the AP site refractory to spontaneous β-elimination or cleavage by AP endonucleases and AP lyases. HMCES thereby prevents DSB formation at AP sites encountered during replication and promotes cellular resistance to a variety of chemical agents and enzymatic activities that directly or indirectly induce AP site formation.

**Figure 1.**
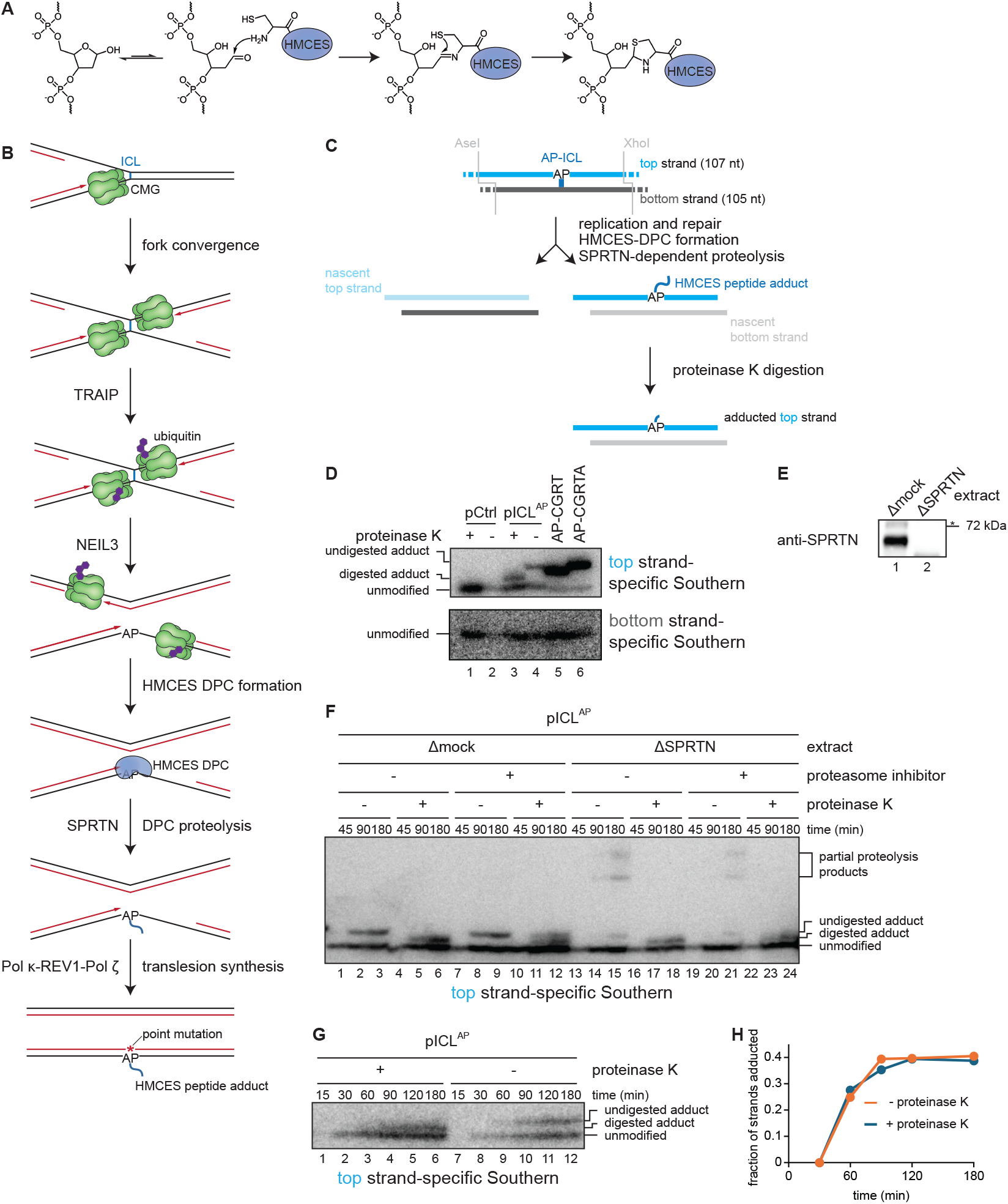
HMCES-DPC processing during ICL repair. **A.** Mechanism of HMCES-DPC formation. **B.** Model of replication-coupled AP-ICL repair by the NEIL3 glycosylase. Green, CMG; black, parental DNA strands; red, nascent DNA strands; purple, ubiquitin; blue, HMCES. **C.** Strategy for resolving repair products. AseI and XhoI digestion allows resolution of the top (107 nt) and bottom (105 nt) strands. AP-ICL unhooking by NEIL3 generates an AP site that cross-links to HMCES and is processed by SPRTN. The resulting adduct can be further digested with proteinase K. **D.** Detection of HMCES adducts by strand-specific Southern blotting. pCtrl or pICL^AP^ was replicated in egg extract supplemented with p97 inhibitor to prevent activation of the Fanconi anemia pathway. Replication products were treated with proteinase K, as indicated, digested with AseI and XhoI, and separated on a denaturing polyacrylamide sequencing gel alongside plasmids containing defined 4 and 5 amino acid peptide-AP site adducts. DNA strands were visualized by Southern blotting with the indicated strand-specific probes. **E.** SPRTN immunodepletion. The extracts used in the replication reactions shown in **F** were blotted for SPRTN. **F.** pICL^AP^ was replicated in the indicated extracts supplemented with the proteasome inhibitor MG262, as indicated. Repair products were resolved and visualized by strand-specific Southern blotting as in **D**. **G.** pICL^AP^ was replicated in extract. At the indicated times, replication products were resolved and visualized as in **D** with and without proteinase K treatment. **H.** The ratio of adducted top strands to unmodified top strands was quantified for the experiment shown in **G**.

Although HMCES is critical for stabilizing ssDNA AP sites during replication, HMCES-DPCs must eventually be removed to allow repair of the underlying DNA lesions. Multiple mechanisms have been reported to contribute to HMCES-DPC removal. HMCES-DPCs strongly accumulate in human cells treated with proteasome inhibitor, implying that the proteasome contributes to the processing and eventual removal of HMCES-DPCs^6,10^. HMCES-DPCs are digested by the replication-associated protease SPRTN in purified systems and cell-free extracts, indicating that SPRTN can also contribute to HMCES-DPC proteolysis^11–13^. Interestingly, the HMCES SRAP domain can catalyze thiazolidine cross-link reversal^12,14,15^. In the context of dsDNA, cross-link reversal leads to HMCES dissociation due to the protein’s reduced affinity for dsDNA, enabling non-proteolytic HMCES-DPC removal and regeneration of an AP site in DNA.

We previously demonstrated that an HMCES-DPC forms as an intermediate during replication-coupled repair of an AP site-derived DNA interstrand cross-link (AP-ICL) in *Xenopus* egg extract (Figure 1B)^11^. An AP-ICL is generated when an AP site reacts with an exocyclic amine of a nucleobase in the opposite strand. This covalent linkage blocks DNA replication, leading to the convergence and stalling of replicative CDC45-MCM2-7-GINS (CMG) helicases at the ICL. Fork convergence promotes the recruitment of the NEIL3 glycosylase, which resolves the ICL by cleaving the *N*-glycosyl bond comprising the cross-link to regenerate an AP site in one strand and an undamaged base in the other^16,17^. The resulting AP site, which is exposed in ssDNA, is cross-linked by HMCES, thereby suppressing AP site cleavage and DSB formation. Ultimately, the HMCES-DPC is digested by the DNA-dependent SPRTN protease and then bypassed by the Pol κ-REV1-Pol ζ TLS polymerase complex. Strikingly, HMCES-DPC formation during ICL repair was observed to introduce a strong nucleotide insertion bias opposite the AP site, suggesting that the HMCES-DPC controls the mutagenicity of the TLS step. However, the mechanism by which an HMCES adduct influences TLS is unknown.

Here, we show that HMCES-DPCs formed during NEIL3-dependent ICL repair are rapidly proteolyzed by SPRTN to yield short, five amino acid peptide adducts. Like DPCs formed by endogenous HMCES, a defined AP site peptide adduct is bypassed by the Pol κ-REV1-Pol ζ TLS polymerase complex, resulting in mutagenic insertion opposite the lesion. Surprisingly, the HMCES peptide adduct does not directly influence insertion of specific nucleotides opposite the AP site. Instead, the identity of the inserted base depends on the template nucleotide 5’ adjacent to the AP site. Our results indicate that the ring-opened thiazolidine adduct promotes template slippage, allowing the nucleobase adjacent to the lesion to template two successive nucleotide incorporations by the TLS polymerase. These findings reveal a previously unrecognized mode of lesion bypass that may be broadly utilized during bypass of DNA adducts that induce AP site ring opening.

## Results

### SPRTN-dependent HMCES-DPC processing produces a 5 amino acid peptide adduct

We previously demonstrated that HMCES-DPCs formed during NEIL3-dependent AP-ICL repair in *Xenopus* egg extracts are digested by the SPRTN protease^11^. However, our results did not distinguish the extent to which SPRTN processes the HMCES-DPC. We therefore set out to determine the length of the HMCES-peptide adduct generated upon SPRTN-dependent proteolysis. Undamaged plasmid (pCtrl) or plasmid containing a site-specific AP-ICL (pICL^AP^) was replicated in *Xenopus* egg extracts, and a restriction fragment encompassing the ICL was resolved on a denaturing polyacrylamide sequencing gel. The top and bottom DNA strands were then detected by strand-specific Southern blotting (Figure 1C). Where indicated, samples were treated with proteinase K to digest the HMCES-DPC to a short peptide adduct that migrates as a single band on sequencing gels^11^. As expected, replication of pCtrl and pICL^AP^ yielded only unmodified bottom strands, confirming that no adducts formed on the undamaged bottom strands (Figure 1D). By contrast, whereas pCtrl replication yielded only unmodified top strands, pICL^AP^ replication produced an additional slower-migrating top strand species, consistent with HMCES-DPC formation during NEIL3-dependent AP-ICL repair (Figure 1D). Strikingly, in the absence of proteinase K digestion, the adducted top strand resolved as a discrete species that migrated more slowly than the proteinase K-digested HMCES adduct. This slower-migrating species co-migrated with a standard prepared by reacting an AP site-containing DNA oligonucleotide with a Cys-Gly-Arg-Thr-Ala (CGRTA) pentapeptide corresponding to the *Xenopus* HMCES N-terminus. We conclude that endogenous proteases process HMCES-DPCs into five amino acid peptide adducts during ICL repair in egg extract.

To determine whether HMCES-DPC processing to a pentapeptide adduct depends on the SPRTN protease, pICL^AP^ was replicated in mock- or SPRTN-depleted egg extract and the adducted top strand was resolved on a denaturing sequencing gel (Figure 1, E and F). After proteinase K treatment, a short HMCES adduct was resolved following replication in either mock- or SPRTN-depleted extract indicating that HMCES-DPCs are produced during ICL repair in both the presence and absence of SPRTN (Figure 1E). However, SPRTN-depletion resulted in disappearance of the pentapeptide adduct that is observed when proteinase K treatment is omitted. We surmise that, in the absence of SPRTN-dependent proteolysis, the unprocessed HMCES-DPC fails to be resolved on the region of the sequencing gel probed by Southern blotting. Notably, addition of the proteasome inhibitor MG262 had no impact on formation of the pentapeptide adduct, implying that the proteasome does not contribute to HMCES-DPC processing in egg extract, consistent with our previous results^11^. In some experiments, we detected low levels of top strand species that may correspond to longer HMCES peptide adducts following replication in SPRTN-depleted extract (Figure 1F). We propose that these species reflect incomplete HMCES-DPC processing by residual SPRTN or an alternative DPC protease. Taken together, these data indicate that SPRTN degrades HMCES-DPCs generated during ICL repair to five amino acid peptide adducts.

We next examined the kinetics of SPRTN-dependent HMCES-DPC proteolysis. Treatment with proteinase K enables resolution of all HMCES-DPCs on sequencing gels, regardless of whether the DPCs have been proteolyzed by SPRTN. In contrast, only HMCES-DPCs that have been already processed to a short adduct are resolved when proteinase K treatment is omitted. We performed a pICL^AP^ replication time course and monitored the accumulation of HMCES adducts with and without proteinase K digestion (Figure 1G). Shortly after initiating replication, the AP site-containing top strand was not detected due to the presence of the AP-ICL, which prevents resolution of the individual DNA strands. As seen previously^11^, total top strand HMCES adducts accumulated more slowly than unmodified top strands in proteinase K-treated samples, indicating that HMCES-DPC formation occurs after a delay following NEIL3-dependent ICL unhooking and AP site generation. Importantly, the appearance of pentapeptide-adducted top strands observed in the absence of proteinase K treatment closely mirrored the appearance of proteinase K-digested adducts (Figure 1, G and H), implying that SPRTN rapidly degrades HMCES upon DPC formation. This finding aligns with a previous report that SPRTN is most active toward DPCs positioned at ssDNA-dsDNA junctions, such as those formed during NEIL3-dependent ICL repair^18^. Taken together, our results indicate that, following NEIL3-dependent ICL unhooking and extension of nascent leading strands up to the resulting AP site, HMCES slowly cross-links to the AP site and is immediately degraded to a pentapeptide adduct by SPRTN.

### HMCES peptide adducts are bypassed by translesion synthesis

Completion of NEIL3-dependent ICL repair in egg extracts requires mutagenic TLS^11,16^. Rapid proteolysis of HMCES-DPCs formed during ICL repair implies that TLS predominantly involves bypass of an HMCES peptide adduct. To directly test whether bypass of an HMCES peptide adduct requires TLS, we constructed plasmid replication templates containing site-specific adducts using AP site-containing oligonucleotides that were reacted with a synthetic CGR tripeptide corresponding to the HMCES N-terminus (Figure 2A and Supplementary Figure S1A). Two plasmids were prepared in which the HMCES tripeptide adduct was positioned in either DNA strand, adjacent to an array of forty-eight LacR binding sites (48x*lacO*). Incubation with LacR results in a barrier that allows only one replication fork to encounter the adduct and in a defined orientation^19^, either in the leading or lagging strand template (pCGR^lead^ or pCGR^lag^). In the absence of LacR, replication forks traveling in either direction could encounter the adduct (see schematic in Figure 2A). Under these conditions, replication of both pCGR^lead^ and pCGR^lag^ resulted in an accumulation of gapped monomeric plasmids that were gradually converted into fully replicated supercoiled plasmids (Supplementary Figure S1B), consistent with a requirement for TLS to bypass the peptide adduct. Similarly, nascent leading strand synthesis stalled one nucleotide upstream of the adduct (−1 stall; Figure 2A) before eventually resuming to produce full-length extension products. No stall products were observed for the nascent leading strands templated by the undamaged parental DNA strand. We also did not detect stall products 20 to 40 nucleotides upstream of the adduct corresponding to CMG helicase stalling at the lesion, implying that the tripeptide adduct does not hinder unwinding by CMG. In the presence of LacR, late replication (θ) intermediates accumulated due to replication fork stalling at the periphery of the *lacO*-LacR barrier (Supplementary Figure S1Β) and only the rightward fork encountered the peptide adduct. Under these conditions, replication of both pCGR^lead^ and pCGR^lag^ again induced pronounced −1 stalling (Figure 2A), indicating that both leading strand synthesis by Pol ε and lagging strand synthesis by Pol δ are blocked by the adduct.

**Figure 2.**
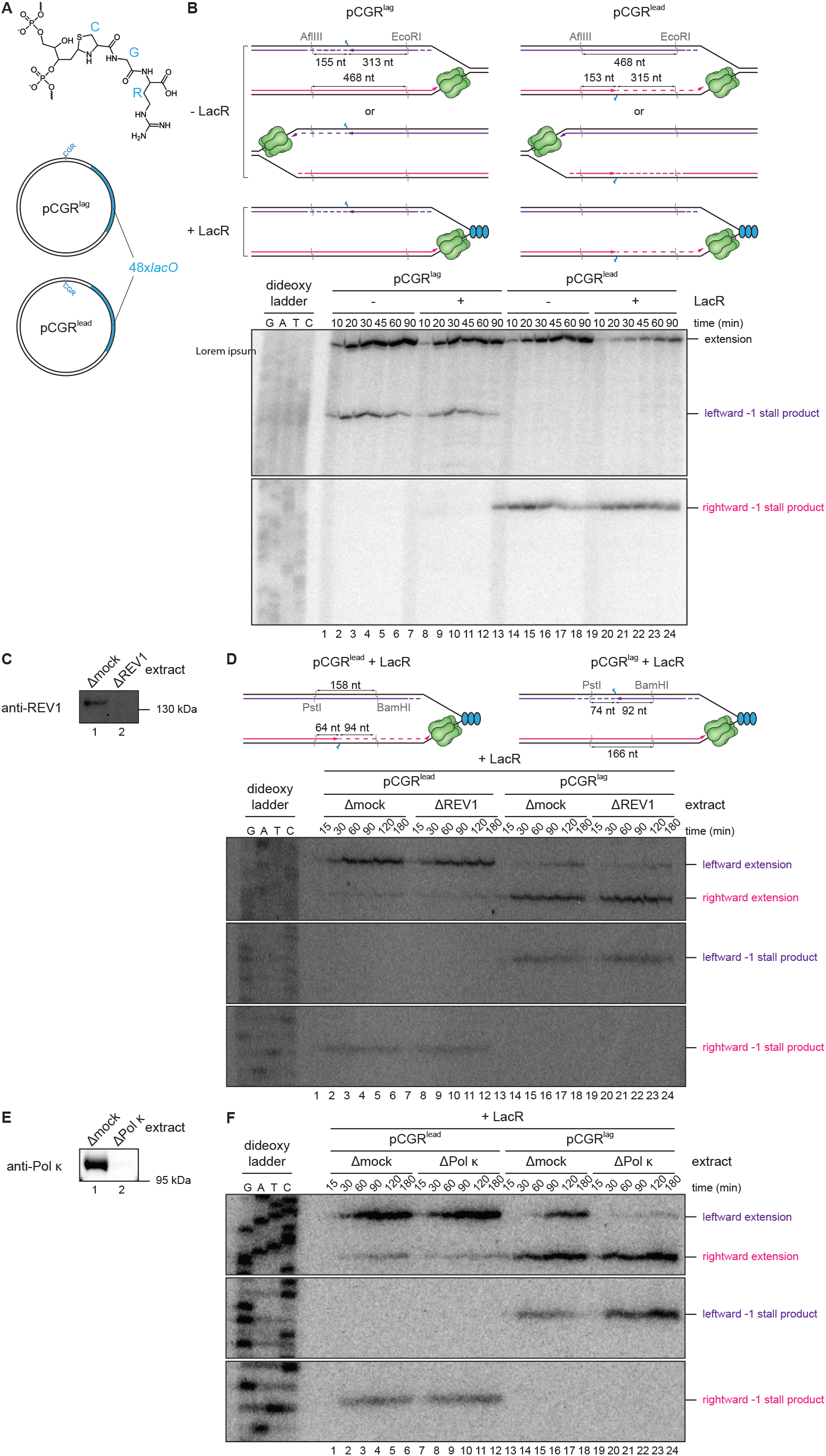
Bypass of an HMCES peptide adduct requires TLS. **A.** Structure of CGR tripeptide AP site adduct. **B.** Top, Schematics of nascent strands generated by replication of pCGR^lag^ and pCGR^lead^ in the absence or presence of LacR. AflIII cuts pCGR^lag^ and pCGR^lead^ 155 nt and 153 nt, respectively, to the left of the peptide adduct. EcoRI cuts pCGR^lag^ and pCGR^lead^ 313 nt and 315 nt, respectively, to the right of the peptide adduct. Bottom, pCGR^lag^ and pCGR^lead^ were replicated with [α-^32^P]dCTP in egg extract supplemented with LacR, as indicated. Replication intermediates were isolated at the indicated times, digested with AflIII and EcoRI, resolved by denaturing polyacrylamide gel electrophoresis, and visualized by autoradiography. Top and bottom panels show sections of the same gel to visualize extension, leftward leading and rightward lagging strands, respectively. **C.** REV1 immunodepletion. The extracts used in the replication reactions shown in **D** were blotted for REV1. **D.** Schematics of nascent strands generated by replication of pCGR^lag^ and pCGR^lead^ in the presence of LacR. PstI cuts pCGR^lag^ 64 nt to the left of the peptide adduct. BamHI cuts pCGR^lead^ 88 nt to the right of the peptide adduct. Bottom, pCGR^lag^ and pCGR^lead^ were replicated in the indicated extracts supplemented with [α-^32^P]dCTP and LacR. Nascent DNA strands were digested with PstI and BamHI and resolved and visualized as in **B**. Top and bottom panels show sections of the same gel to visualize extension, leftward leading and rightward lagging strands, respectively. **E.** Pol κ immunodepletion. The extracts used in the replication reactions shown in **F** were blotted for Pol κ. **F.** pCGR^lag^ and pCGR^lead^ were replicated in the indicated extracts supplemented with [α-^32^P]dCTP and LacR. Nascent DNA strands were digested with PstI and BamHI and resolved and visualized as in **B**. Top and bottom panels show sections of the same gel to visualize extension, leftward leading and rightward lagging strands, respectively.

Despite identical sequences surrounding the leading and lagging strand template adducts, bypass of the leading strand adduct was markedly slower than bypass of the lagging strand adduct. We wondered if different TLS polymerases are used to bypass HMCES peptide adducts when the lesion is encountered in the leading strand or lagging strand templates. During ICL repair, the requirement for fork convergence to activate ICL unhooking by NEIL3 dictates that all encounters with HMCES-DPCs occur on the leading strand template (Figure 1A). In this case, TLS past the leading strand HMCES-DPC requires a polymerase complex comprising Pol κ, REV1, and Pol ζ, with Pol ζ thought to function as the catalytic subunit^20^. Replication of pCGR^lead^ in REV1- or Pol κ-depleted extracts supplemented with LacR, which prevented fork convergence (Supplementary Figure S1, C and D), resulted in a persistence of leading strand stall products and a corresponding decrease in the formation of full-length extension products (Figure 2, B-E), indicating that bypass of leading strand HMCES adducts depends on REV1 and Pol κ. Importantly, depletion of REV1 or Pol κ also strongly inhibited conversion of stall products into full-length extension products on the lagging strand during pCGR^lag^ replication (Figure 2, B-E and Supplementary Figure S1, C and D). Our data are consistent with use of the same Pol κ-REV1-Pol ζ complex to bypass HMCES adducts present on both the leading and lagging strand templates. This finding aligns with a recent report that bypass of an AP site proceeds by the same REV1-dependent mechanism during leading and lagging strand TLS in egg extract^21^. We speculate that differences in the rate of adduct bypass during leading and lagging strand synthesis may reflect differences in the rate of polymerase exchange on the two strands. Alternatively, differences in template structure caused by helicase-polymerase uncoupling when a leading strand template lesion is present may contribute to slower leading strand TLS.

### The mutagenicity of HMCES bypass depends on AP site ring opening

During NEIL3-dependent ICL repair in egg extract, HMCES-DPC formation alters the mutagenicity of TLS past an AP site^11^. In the absence of HMCES-DPC formation, TLS follows the “A-rule”^22^, resulting in preferential insertion of deoxyadenosine (dA) opposite the unprotected AP site. We previously showed that HMCES-DPC formation suppressed dA insertions opposite the adduct, giving rise to a corresponding increase in deoxyguanosine (dG) insertions. This led us to speculate that an HMCES peptide adduct might directly template dG insertion opposite an AP site. Such a mechanism would be analogous to dC insertion mediated by REV1, which uses a conserved arginine to direct dCTP incorporation opposite an AP site^23^. To determine whether an HMCES tripeptide adduct is sufficient to produce the mutation spectrum observed during ICL repair, we replicated pCGR in egg extract and analyzed the repair products using next-generation sequencing (Figure 3A and Supplementary Figure S2, A and B). Similar to pICL^AP^ replication products, pCGR replication products were generally free of insertions or deletions and contained primarily dG and dA insertions opposite the adduct (Figure 3B), indicating that the tripeptide adduct recapitulates the mutagenicity of HMCES-DPC bypass. dG was the most common nucleotide inserted regardless of whether the adduct was encountered on the leading strand template or lagging strand template (Figure 3C and Supplementary Figure S2, C and D), providing additional evidence that the same TLS mechanism promotes bypass of both leading and lagging strand HMCES-DPCs.

**Figure 3.**
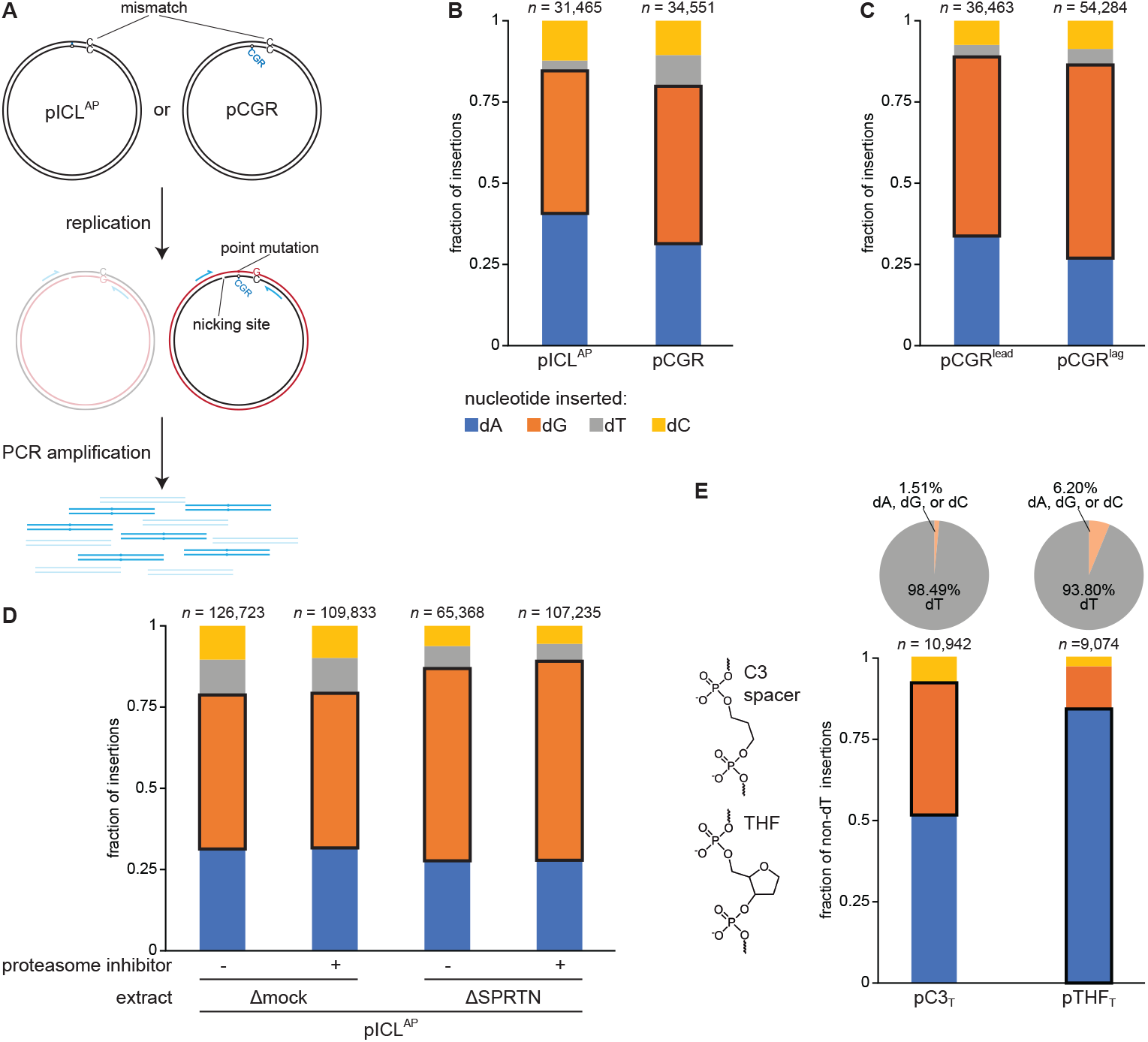
A ring-open AP site recapitulates the mutation spectrum produced by HMCES during ICL repair. **A.** Schematic describing method for sequencing nascent strands produced upon AP site bypass. Incorporation of a C-C mismatch distinguishes DNA strands templated by the undamaged bottom strand from those templated by the adducted top strand. Nicking with Nt.BstNBI ensures that adducted parental DNA strands are not amplified during library preparation. **B.** pICL^AP^ and pCGR^lag^ were replicated in egg extract, and sequencing libraries were prepared from replication products as described in A. The fraction of reads corresponding to insertion of a given nucleotide opposite the AP site is plotted for each plasmid. *n*, number of nascent strand reads obtained for each plasmid. Next-generation sequencing read counts can be found in Table X. **C.** pCGR^lag^ and pCGR^lead^ were replicated in egg extract supplemented with LacR and replication products were sequenced as in **B**. **D.** pICL^AP^ was replicated in the indicated extracts supplemented with proteasome inhibitor MG262, as indicated, and replication products were sequenced as in **B**. **E.** Left, structures of C3 spacer and THF AP site analogs. pC3_T_ and pTHF_T_ were replicated in egg extract and replication products were sequenced as in **B**. Top, pie chart depicting the fraction of reads corresponding to dT and non-dT insertions opposite the AP site analog. Bottom, the fractions of non-dT reads corresponding to insertion of dA, dG, and dC opposite the AP site analog is plotted. *n*, number of nascent strand non-dT insertion reads.

We next tested whether proteolysis of the HMCES-DPC is required for the mutation spectrum produced during NEIL3-dependent ICL repair. pICL^AP^ was replicated in mock- or SPRTN-depleted extract supplemented with proteasome inhibitor, and the replication products were sequenced (Figure 3D and Supplementary Figure S2, E and F). Once again, upon replication in mock-depleted extract, dG insertion opposite the AP site occurred most frequently (47.55% of insertions; Figure 3C). Importantly, the frequency of dG insertion slightly increased in SPRTN-depleted extract (to 59.33% of insertions), indicating that SPRTN-dependent proteolysis of the HMCES-DPC is not required to produce the dG insertion preference during NEIL3-dependent ICL repair. Proteasome inhibition with MG262 did not affect the mutation spectrum observed in mock- or SPRTN-depleted extracts, consistent with the lack of a role for the proteasome during HMCES-DPC processing in egg extract.

To determine the minimum AP site structure necessary to confer a bias for dG insertion during bypass, we examined the mutagenicity of a three carbon (C3) spacer (Figure 3E). The C3 spacer mimics the ring-open form of the AP site generated upon formation of the HMCES thiazolidine linkage. Plasmid containing a site specific C3 spacer (pC3_A_) in a sequence context identical to that of the pCGR plasmids was replicated in egg extract and the replication products were sequenced (Supplementary Figure S2, G and H). In contrast to the pCGR replication products, the overwhelming majority of pC3_A_ replication products (98.54%) corresponded to dA insertion opposite the C3 spacer position (Supplementary Figure SI). However, dA also corresponds to the base that would be observed following error-free repair of the C3 spacer-containing strand. APE1 endonuclease can efficiently incise 5’ of a C3 spacer to initiate BER^24^, suggesting that the C3 spacer might be efficiently removed by BER in egg extract. Indeed, while pC3_A_ was initially sensitive to nicking by recombinant APE1, this sensitivity was lost following incubation in egg extract, consistent with repair (Supplementary Figure S2J). To distinguish dA insertions that result from TLS from those that result from BER, we prepared a C3 spacer plasmid with dT positioned opposite the lesion (pC3_T_). As expected, 98.48% of sequenced pC3_T_ replication products corresponded to dT insertion opposite the lesion, consistent with efficient repair of the C3 spacer (Figure 3E and Supplementary Figure 2, K and L). The remaining 1.51% of reads produced by pC3_T_ replication corresponded to dA, dG, or dC insertion opposite the C3 spacer. Although this fraction of reads represents a small proportion of the total reads derived from replication on the strand containing the C3 spacer, it is much higher than the 0.10% sequencing error rate observed across the entire sequencing library amplicon. We therefore infer that the non-dT insertion reads reflect bypass of unrepaired C3 spacers. Interestingly, dA and dG comprised 51.49% and 40.30%, respectively, of the non-dT insertions (Figure 3E). This relative rate of dA and dG insertions paralleled the rates of dA and dG insertion observed for pICL and pCGR plasmids, implying that a common mechanism produces the observed dG insertion biases in all three cases.

For comparison, we also sequenced the replication products of a plasmid containing a tetrahydrofuran (THF) AP site analog that mimics the hemiacetal form of an AP site but cannot tautomerize to the ring-opened aldehyde form that is cross-linked by HMCES (pTHF_T_; Figure 3E and Supplementary Figure 2, K and L). Once again, the overwhelming majority of reads corresponded to dT insertion opposite the lesion (93.80%; Figure 3E), consistent with efficient THF removal by BER (Supplementary Figure 2J). In contrast to pC3T replication products, the remaining pTHF_T_ replication products contained a much greater proportion of dA insertions (83.99%) than dG insertions (12.94%). This preference for dA insertion during TLS past THF is similar to the preference for dA insertion observed during bypass of an unprotected AP site generated during NEIL3-dependent ICL repair in HMCES-depleted extract^11^. Combined, these data suggest that it is not the formation of an HMCES-DPC *per se* that favors dG insertion opposite an AP site but rather the formation of a stable ring-opened form of the AP site.

### The template nucleotide 5’ of an HMCES-peptide adduct influences insertion opposite the lesion

In considering how a ring-open AP site could exhibit a mutation spectrum distinct from that of ring-closed AP site, we realized that a dC nucleotide was positioned immediately 5’ of the DNA lesion in each of our previous experiments (Figure 4A). This suggested a model in which the ring-open AP site accommodates a shift in the template strand register that allows the adjacent nucleotide to template insertion opposite the lesion. Following this insertion event, the template strand would revert to its original register, allowing the position 5’ of the AP site to template a second round of nascent strand extension. Ultimately, this template slippage would result in production of a full-length nascent strand. To test this model, we first prepared an HMCES tripeptide-containing plasmid with dG 5’ of the lesion (pCGR^5’dG^; Figure 4A and Supplementary Figure S1A). The plasmid was otherwise identical to pCGR^lag^ (referred to here as pCGR^5’dC^), which has dC 5’ of the adduct. We then replicated the plasmid in egg extract and sequenced the replication products (Figure 4B and Supplementary Figure S3, A and B). When dC was positioned 5’ of the tripeptide adduct, dG insertions opposite the adduct occurred most frequently (43.85%), while dC insertions occurred infrequently (12.28%). However, when dG was positioned 5’ of the tripeptide adduct, dG insertions opposite the adduct were dramatically reduced (5.95%), while dC insertions dramatically increased (35.68%). This result implies that the base inserted opposite the tripeptide adduct is templated by the nucleotide 5’ of the adduct.

**Figure 4.**
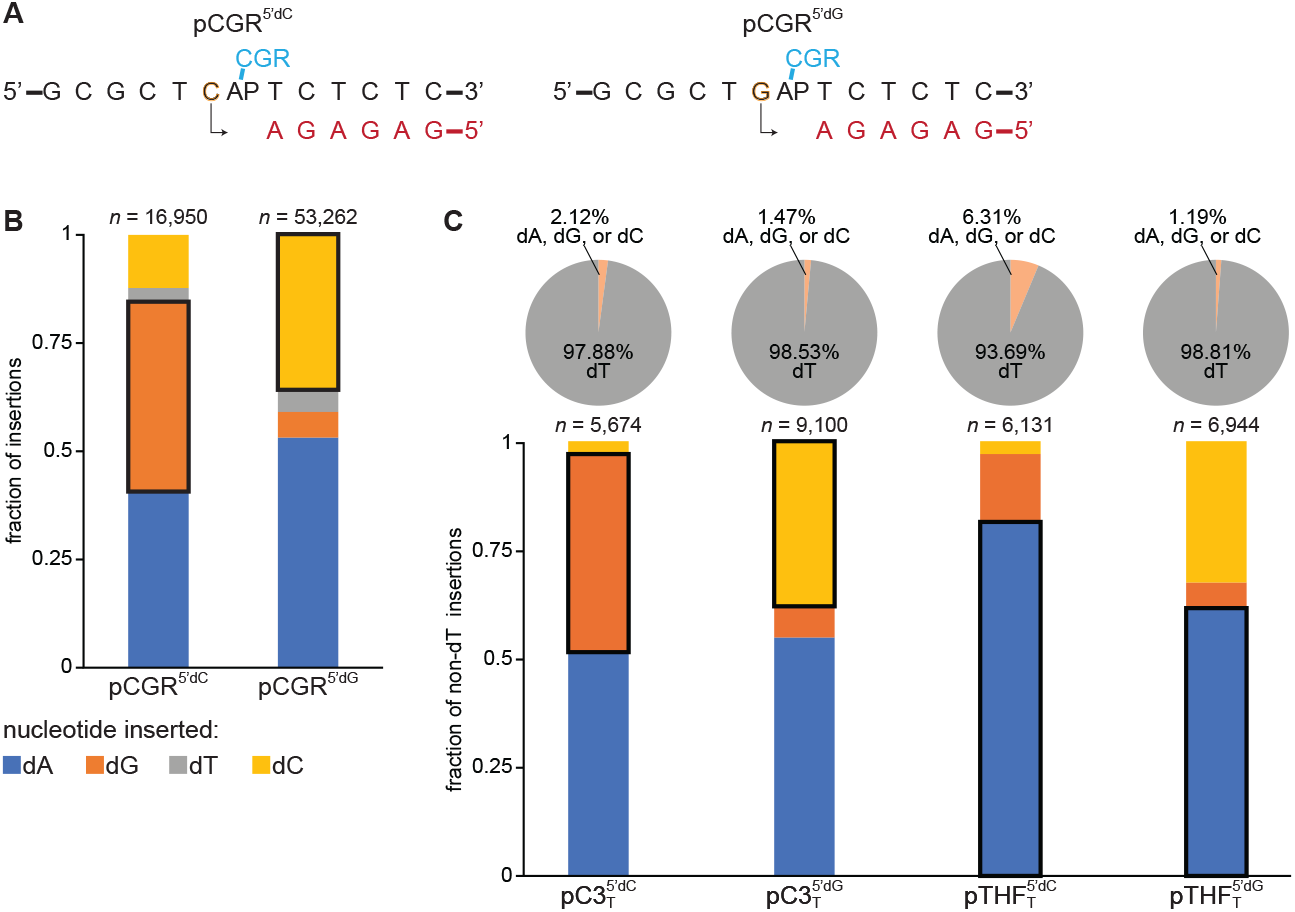
The nucleotide 5’ of the HMCES-adducted AP site templates insertion. A. Schematic of pCGR^5’dC^ and pCGR^5’dG^ replication. Nascent strands are shown in red. The base 5’ of the adducted AP site is outlined in orange. B. pCGR^5’dC^ and pCGR^5’dG^ were replicated in egg extract supplemented and replication products were sequenced as in Figure 3B. The fractions of reads corresponding to insertion of dA, dG, dT, and dC opposite the AP site analog is plotted. *n*, number of nascent strand reads obtained for each plasmid. C. pC3T and pTHFT plasmids with the indicated template nucleotides 5’ of the AP site analog were replicated in egg extract and replication products were sequenced as in Figure 3B. Top, pie chart depicting the fraction of reads corresponding to dT and non-dT insertions opposite the AP site analog. Bottom, the fractions of non-dT reads corresponding to insertion of dA, dG, and dC opposite the AP site analog is plotted. *n*, number of nascent strand non-dT insertion reads.

We also examined the effect of the 5’ template nucleotide during bypass of C3 spacer and THF AP site analogs (Figure 4C and Supplementary Figure S3, C and D). As seen in Fig. 3E, replication of pC3_T_ with a 5’ template dC resulted in similar rates of dA and dG insertion (51.55% and 45.36% of non-dT insertions, respectively) with relatively few dC insertions (3.09%), whereas replication of pTHF_T_ with a 5’ template dC resulted greater proportions of dA than dG insertions (81.29% and 15.81% of non-dT insertions, respectively). Consistent with a role for the template 5’ nucleotide in directing insertion opposite the lesion, replication of pC3_T_ with a template dG 5’ of the lesion produced a reduction in dG insertions (7.32%) and a corresponding increase in dC insertions (37.80%). Surprisingly, a notable increase in the frequency of dC insertions (32.35%) was also observed for THF plasmid containing dG 5’ of the lesion, although dA insertions were still the most prevalent (61.76%). Thus, a ring-closed AP site may also support template slippage during TLS, at least in some sequence contexts. In aggregate, our results indicate that the template nucleotide 5’ of a ring-opened HMCES adduct can direct nascent strand base insertion at the position corresponding to the lesion.

### The template 5’ nucleotide directs insertion during bypass of endogenous HMCES-DPCs

We next comprehensively evaluated how the nucleotide 5’ of the AP site influences the mutation spectrum generated during NEIL3-dependent AP-ICL repair. AP-ICL-containing plasmids with each of the four nucleotides immediately 5’ to the AP site were replicated in parallel and the replication products were sequenced (Figure 5A and Supplementary Figure S5, A and B). In all cases, dA insertion represented the most common or second most common class of replication product, accounting for 31.97-55.51% of reads derived from bypass of the AP site. This result is consistent with the established preference for dA insertion during AP site bypass observed in our previous experiments^11^. However, the base inserted opposite the AP site was otherwise correlated with nucleotide 5’ of the AP site: the frequency of dG insertions was highest (49.61%) when dC was present in the template strand, the frequency of dC insertions was highest (25.69%) when dG was present, the frequency of dA insertions was highest (55.51%) when dT was present and the frequency of dT insertions was highest (25.51%) when dA was present. Importantly, the strength of the correlation between the identity of the inserted base and the identity of the base 5’ of the AP site varied across the four plasmids. This could reflect differences in the ability of HMCES to cross-link the AP site in different sequence contexts or differences in the ability of each template nucleotide to be accommodated within the TLS polymerase active site in the shifted register. In any case, these results indicate that bypass of an endogenous HMCES-DPC can involve a template slippage mechanism.

**Figure 5.**
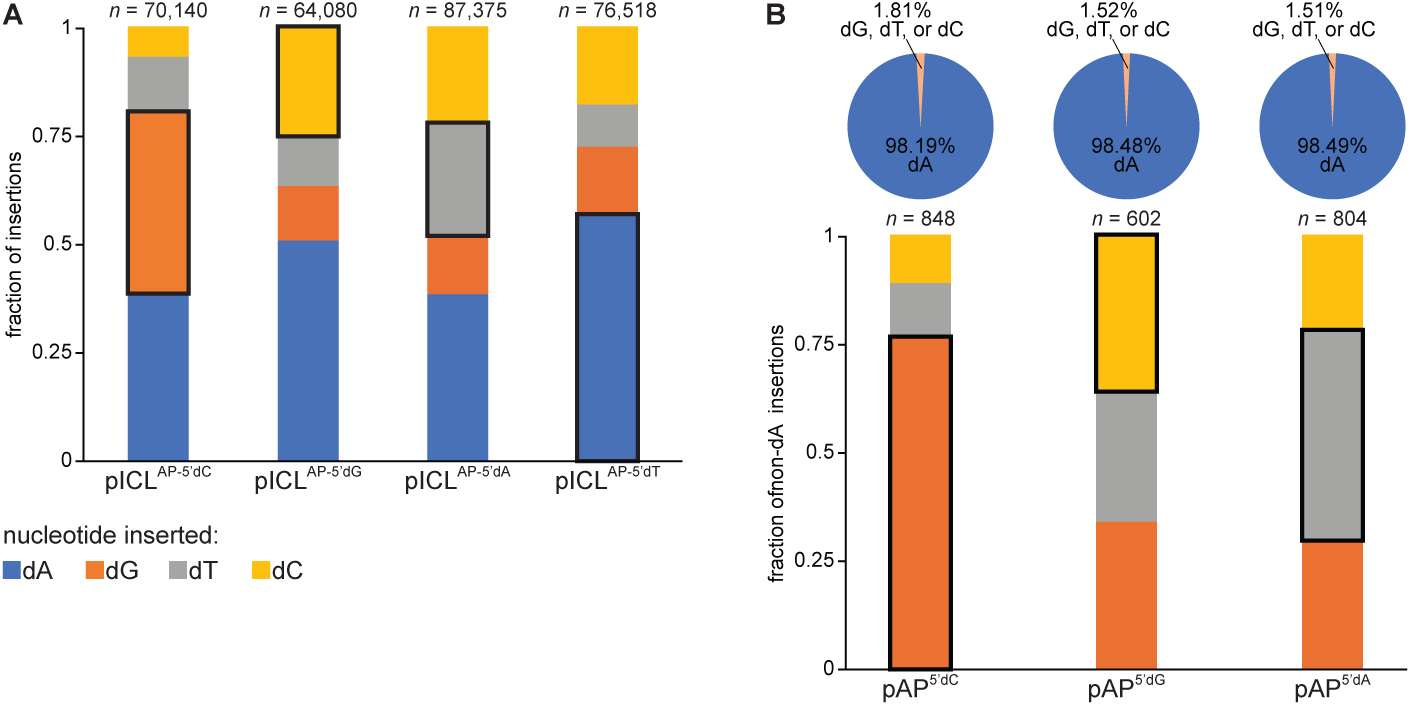
HMCES-DPCs promote template slippage. **A.** pICL^AP^ plasmids with the indicated nucleotide 5’ of the cross-linked AP site were replicated in egg extract and replication products were sequenced as in Figure 3B. The fractions of reads corresponding to insertion of dA, dG, dT and dC opposite the AP site is plotted. *n*, number of nascent strand reads obtained for each plasmid. **B.** pAP plasmids with the indicated nucleotide 5’ of the unprotected AP site were replicated in egg extract and replication products were sequenced as in Figure 3B. Top, pie chart depicting the fraction of reads corresponding to dA and non-dA insertions opposite the AP site. Bottom, the fractions of non-dA reads corresponding to insertion of dG, dC, and dT opposite the AP site is plotted. *n*, number of nascent strand non-dA insertion reads.

The majority of HMCES-DPCs formed during cellular DNA replication are most likely produced when a replication fork unwinds DNA containing a pre-existing AP site. We therefore tested whether the template 5’ nucleotide influences the mutation spectrum produced when a replication fork encounters a preexisting AP site. Accordingly, we prepared different AP site-containing plasmids with dC, dG, or dA immediately 5’ of the AP site (pAP^5’dC^, pAP^5’dG^, and pAP^5’dA^, respectively). We did not analyze the effect of positioning dT upstream of the AP site since the effects of the 5’ dT on insertion could not be distinguished from the effects of AP site repair. The plasmids were replicated in egg extract, and the products were sequenced (Figure 5B and Supplementary Figure 4, C and D). As seen for pC3 replication products, the pAP replication products almost exclusively reflected dA insertions at the AP site, indicative of efficient error-free AP site repair prior to arrival of a replication fork. However, the remaining 1-2% of replication products corresponded to dG, dC, or dT insertions at the AP site. Once again, this minor fraction of reads with dG, dC, or dT at the AP site was much higher than the sequencing error rate for this experiment (0.15%), implying that the reads containing dG, dC, or dT are derived from replication forks that encounter unrepaired AP sites. Once again, the frequency of dG, dC, and dT insertion correlated with the nucleotide 5’ of the AP site: the frequency of dG insertions was highest (76.60%) when dC was present in the template strand, the frequency of dC insertions was highest (35.96%) when dG was present, and the frequency of dT insertions was highest (48.24%) when dA was present. Combined, these data suggest that mutagenic bypass of HMCES-DPCs formed at preexisting AP sites is also directed by the template nucleotide 5’ of the AP site.

### AP site ring opening is associated with increased template flexibility

Our results suggest that HMCES-DPC formation and attendant AP site ring opening permits the template nucleotide 5’ of the AP site to displace the lesion and occupy the TLS polymerase active site. To explore these predicted template dynamics, we conducted molecular dynamics (MD) simulations of a 14-nucleotide (nt) AP site- or C3-spacer-containing DNA template strand in complex with a 10-nt nascent DNA strand primer and the yeast REV3 subunit of Pol ζ. A cryogenic electron microscopy (cryo-EM) structure of yeast Pol ζ holoenzyme bound to a primed 3’ junction (PDB 7S0T) was selected as the basis for simulating template dynamics^25^. For reference, the yeast and *Xenopus* REV3 polymerase domains are 43% identical (62% similar). The DNA template strand position templating incoming dNTP incorporation (T_0_) was substituted with either a ring-closed AP site or a C3 spacer. A C3 spacer, which recapitulates the mutagenicity of an HMCES adduct (Fig. 3E), was used in place of a thiazolidine AP site adduct to simplify model generation (Supplementary Figure 5A).

Three independent 500 ns MD simulations were performed for each system (Supplemental Movies S1-6). For all simulations, the average REV3 backbone Cα atom root mean square deviations (RMSDs) relative to the initial structure rapidly converged to ∼3 Å (Supplementary Figure S5B), indicating that the overall REV3-DNA complex assemblies remained stable and allowing us to compare the local structural dynamics at the lesion. Tracking the position of the AP site 5’ phosphate indicated that the AP site remained relatively constrained across the three simulations (Figure 6, A and C). The template 5’ nucleotide (T_-1_) also maintained a position similar to that observed in the starting model, with the base everted from the REV3 active site and unstacked with the nascent DNA duplex. By contrast, the C3 spacer 5’ phosphate exhibited more dynamical conformational sampling (Figure 6, B and C). Consistently, in each simulation, the RMSD of the lesion 5’ phosphorous atom relative to the starting model was greater for the C3 spacer template (7.52 ± 1.44 Å, 4.83 ± 1.43 Å, 5.34 ± 0.88 Å; Supplementary Figure 5C) than for the AP site template (1.82 ± 0.72 Å, 2.07 ± 0.79 Å, 2.27 ± 0.68 Å). Torsion angle distributions further indicate that the C3 spacer exhibited increased local flexibility for both the lesion and T_-1_ nucleotide (Figure 6, D-F). Whereas the AP site backbone was largely confined to a narrow set of torsion angles in each trajectory, the C3 spacer backbone sampled multiple torsional states across all trajectories. These observations suggest that the ring-opened form of an AP site can access alternative backbone conformations that could potentially facilitate lesion displacement. In one C3 spacer trajectory, the T_-1_ nucleotide reoriented toward the incoming dATP and REV3 palm region (Figure 6B). We note that this movement was not driven by interactions between the T_-1_ base and the incoming dATP, as no direct base pairing was observed during the simulation. Instead, the T_-1_ C formed stable interactions with REV3 in the form of hydrogen bonds to T916 and T917, and a π-π stacking interaction with W808 (Supplementary Figure S5, D and E). It should be noted, however, that these residues are not well conserved between yeast and vertebrate REV3 orthologs. Nevertheless, our results provide insight as to how increased template dynamics might enable the REV3 active site to sample the 5’ template base. Together, these MD simulations support a model in which AP site ring opening increases the local conformational flexibility of the template backbone, thereby facilitating templated nucleotide incorporation using the nucleotide 5’ of the lesion.

**Figure 6.**
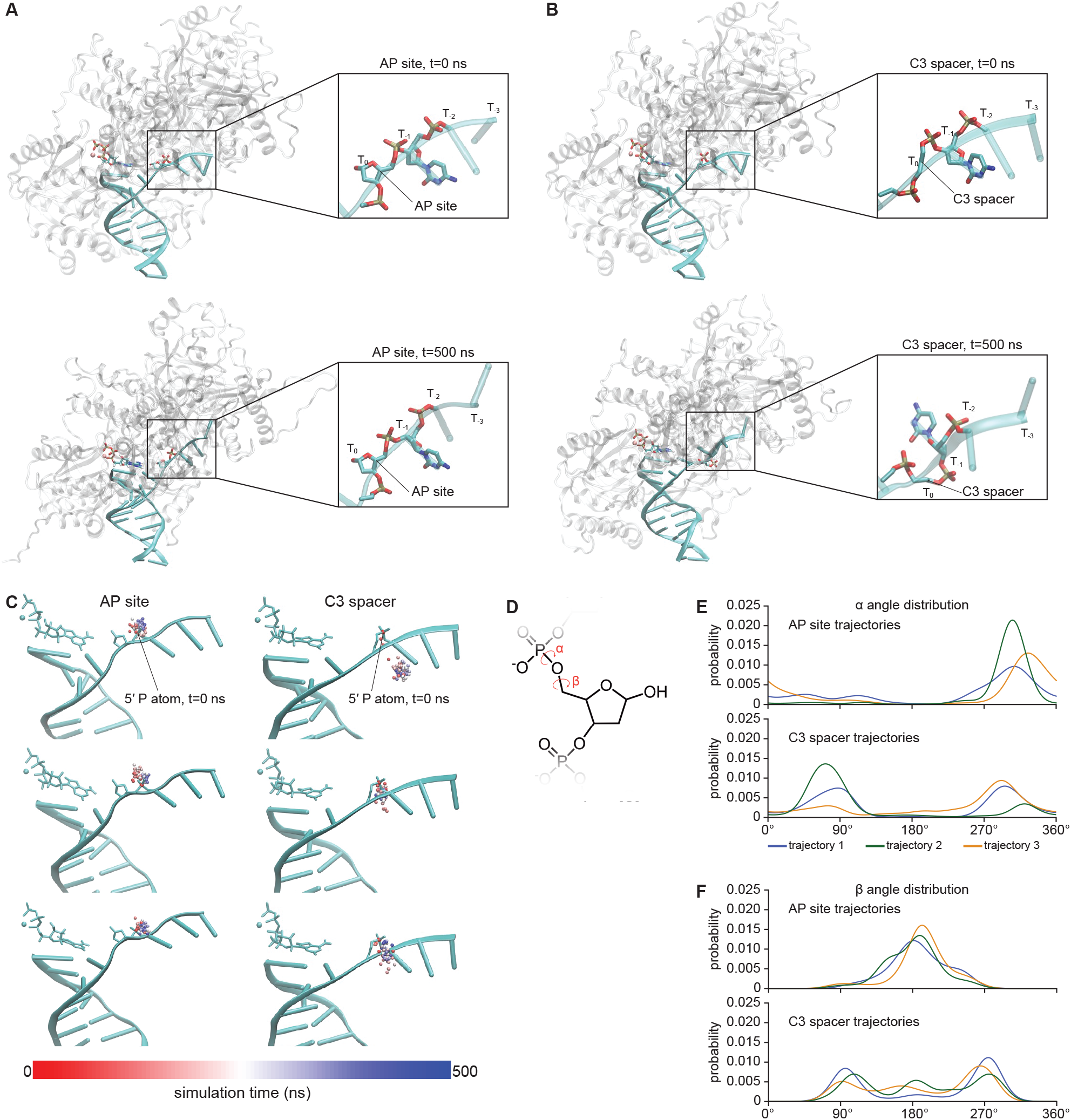
Simulation of AP site and C3 spacer template dynamics. **A** and **B.** Top, Representative starting structures of REV3-AP site DNA (**A**) and REV3-C3 spacer DNA (**B**) complexes used for MD simulation. Bottom, representative structures of REV3-DNA complexes obtained at the end of 500 ns MD simulations. Insets show the local template conformation of the AP site or C3 spacer and 5’ nucleotides in each structure. Gray, yeast REV3 subunit; cyan, DNA duplex. **C.** Positions of the lesion residue 5’ phosphorus (P) atoms are shown for the indicated times during each 500 ns simulation. Left, AP site system. Right, C3 spacer system. Sampled P atom positions are depicted as spheres overlayed on the initial structures (t = 0 ns) of the DNA duplex, dATP and Mg^2+^ ion. Spheres depicting P atoms are color-coded to reflect simulation time. **D.** Schematic depicting DNA backbone α and β dihedral angles. Torsion angle α is the backbone dihedral defined by O3’_(i-1)_–P_(i)_–O5’_(i)_–C5’_(i)_, where i refers to the lesion position. Torsion angle β is the backbone dihedral defined by P_(i)_–O5’_(i)_–C5’_(i)_–C4’_(i)_. **E and F.** Periodic kernel density estimates of the α (**E**) and β (**F**) dihedral angle probability distributions are shown 500 ns simulation. Top, angle distributions for AP site trajectories. Bottom, angle distributions for C3 spacer trajectories.

**Figure 7.**
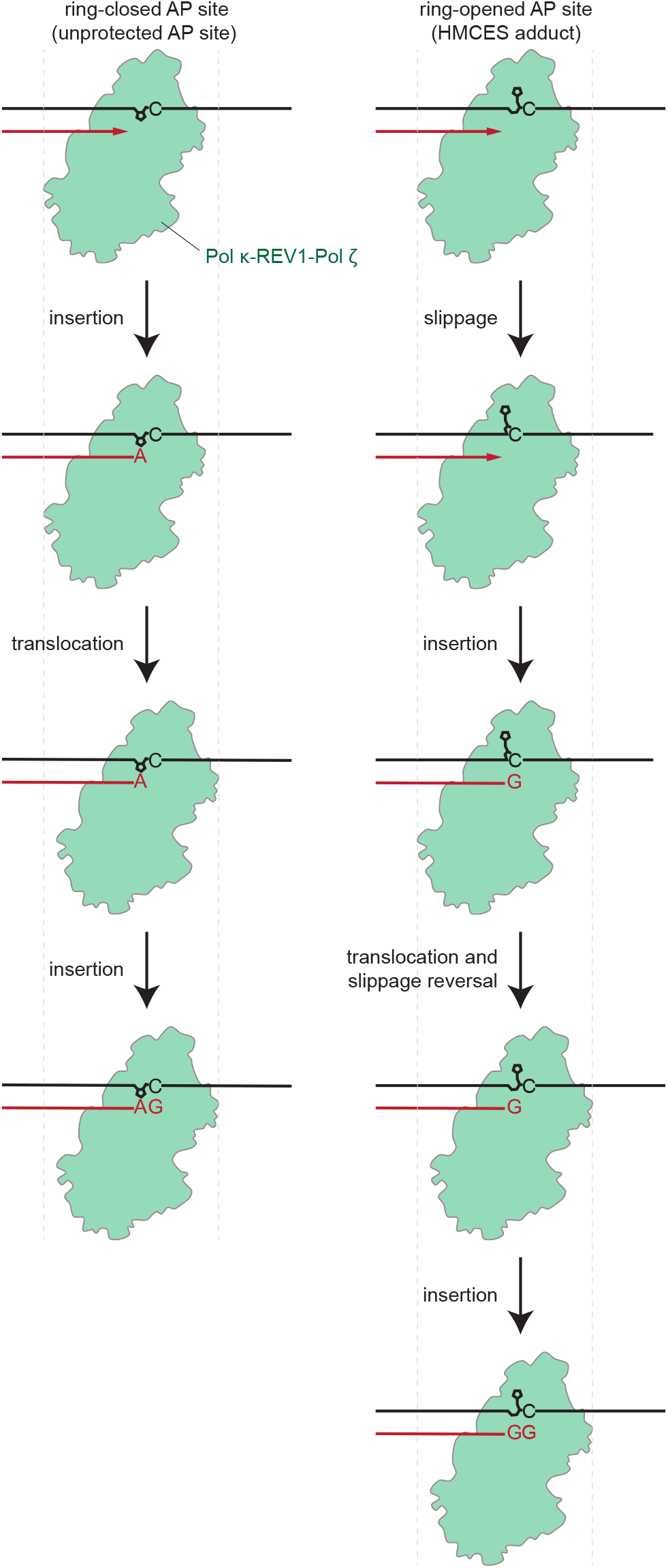
Model for template slipping during HMCES peptide adduct bypass. Left, unprotected ring-closed AP site bypass is biased toward insertion of dA opposite the non-instructive AP site. Right, HMCES adducted ring-opened AP site bypass involves template slippage that enables the nucleotide 5’ of the adduct to template two rounds of insertion. See main text for details. Green, Pol κ-REV1-Pol ζ; black, parental DNA strands; red, nascent DNA strands. Hemiacetal AP sites and thiazolidines are depicted as five member rings.

## Discussion

HMCES-DPC formation plays a critical role in suppressing AP site-induced genome instability during DNA replication. However, the interplay between HMCES-DPC processing and AP site bypass is not well understood. In different contexts, HMCES-DPCs resolution can involve DPC reversal or proteolysis and may be bypassed using template switching or TLS^6,10–12,14,15^. In egg extracts, HMCES-DPCs formed during NEIL3-dependent ICL repair are degraded by SPRTN and bypassed by TLS, introducing a distinct spectrum of mutations. Here, we show that SPRTN digests HMCES-DPC to a five amino acid peptide adduct that is sufficient to block nascent strand extension by replicative polymerases. We also find that the pattern of mutations introduced during TLS past an HMCES-adducted AP site results from stabilization of the ring-opened form of the AP site and is independent of the length of the HMCES peptide adduct. Instead, our results indicate that AP site ring-opening promotes a bypass mechanism involving template slippage (Figure 6). In this model, the ring-opened AP site adopts an alternative backbone conformation that enables the template base 5’ of the AP site to access the polymerase active site and direct insertion. Subsequent translocation by the polymerase is accompanied by slippage reversal that restores the original template register and allows the same base to template a second insertion event, resulting in a nascent strand that contains a point mutation but is free of insertions or deletions. By contrast, the ring closed form of the unprotected AP site is not expected to accommodate template slippage and the TLS polymerase inserts dA opposite the uninstructive lesion, according to the A rule. In support of this model, our MD simulations indicate that a ring-opened AP site analog exhibits considerably more conformational dynamics than a ring-closed AP site.

In egg extracts, TLS past an HMCES peptide adduct relies on the Pol κ-REV1-Pol ζ polymerase complex^20^. However, biochemical evidence indicates that multiple polymerases including Pol η, Pol κ, and Pol δ can extend a primer past a short HMCES peptide adduct, albeit inefficiently^15^. It will be important to determine whether ring-opened HMCES adduct bypass by other DNA polymerases involves template slippage. Interestingly, when extending primers intended to mimic insertion of each of the four nucleotides opposite an HMCES adduct, Pol δ was reported to introduce a single nucleotide deletion only when the primer 3’ terminal nucleotide was complementary to the template nucleotide 5’ of the adduct^15^. This result implies that Pol δ can extend a primer that has effectively bypassed the adduct by realigning to the 5’ template nucleotide. In contrast, deletion products were not observed during extension of the same primers annealed to a template that contained a THF AP site analog. Although this situation is distinct from TLS past an HMCES-DPC in egg extract, in which an iterative forward and reverse slippage produces full-length extension products, it is consistent with a model in which HMCES-DPC induced ring-opening increases the conformational flexibility of the DNA template.

Our data indicating that the mutagenicity of HMCES-DPC bypass is insensitive to the extent of DPC proteolysis is somewhat surprising given the extensive set of contacts between an intact HMCES-DPC and ssDNA. However, this finding is rationalized by previous work demonstrating that the FANCJ helicase stimulates both HMCES-DPC proteolysis by SPRTN and HMCES-DPC bypass by REV1-Pol ζ^13^. FANCJ is a 5’ to 3’ DNA helicase that has been implicated in a variety of genome maintenance pathways including ICL repair, DPC repair, and G quadruplex resolution^13,26,27^. During DPC repair, FANCJ is thought to load onto ssDNA downstream of a DPC and translocate back toward the adduct, unfolding it. This unfolding renders the DPC more susceptible to proteolysis and, in the absence of proteolysis, bypass by TLS polymerases. Thus, during ICL repair in the absence of SPRTN, FANCJ may unfold the HMCES SRAP-domain, disrupting its contacts with DNA and rendering the DPC functionally equivalent to a peptide adduct.

A key question raised by our work is the extent to which ring-opened forms of AP sites exist in cells. In principle, any protein can form a DPC by reacting via a lysine residue with the aldehyde form of an AP site. Beyond HMCES, numerous eukaryotic DNA repair factors react with AP sites to form ring-open Schiff base intermediates. These factors include Pol β, OGG1, NEIL1/2/3, NTHL1, APE1, PARP1, Ku, and histones^28–32^. Highly abundant nuclear polyamines such as spermine and spermidine have also been shown to produce imines at AP sites. However, in all these cases, Schiff base formation is associated with rapid β elimination and strand scission. Thus, while it has been suggested that thousands of DPCs may form at AP sites each day in a human cell^28^, few Schiff base DPCs are likely to be present at steady state in uncleaved DNA that can template replication. Interestingly, endogenous reducing agents such as ascorbic acid and glutathione have been reported to reduce Schiff bases formed between dG and acetaldehyde^33^. It is therefore possible that some AP site DPCs may become stabilized by endogenous reducing agents. Beyond endogenous proteins and metabolites, small-molecule drugs can also trap AP sites in ring-opened forms. Alkoxyamines, which are used as DNA repair inhibitors in combination with other DNA damaging agents, react with AP sites via a Schiff base linkage to produce cleavage resistant adducts^34^. Similarly, the antihypertensive agent hydralazine reacts with AP sites to form a Schiff base intermediate that undergoes spontaneous oxidative cyclization to produce a ring-opened triazolo[3,4-*a*]phthalazine adduct^35,36^. These therapeutics might therefore produce mutational spectrums resembling that produced by HMCES-DPCs. Additional studies are required to determine the cellular prevalence of ring-open AP site derivatives and their impact on mutagenesis.

Finally, it will also be important to evaluate whether template slippage at AP sites plays a role in shaping or maintaining nucleotide frequencies across different species. Observed dinucleotide frequencies often diverge from expectation in many species. CpG and TpA are known to be underrepresented among vertebrate genomes due to their effects on DNA methylation and RNA stability, respectively^37^. Conversely, many species exhibit overrepresentation of homodinucleotides, especially AA/TT, a fact that has been attributed to template slippage^38^. TLS past HMCES adducts may therefore provide a source of new homodinucleotides. Alternatively, template slippage at AP sites may represent a conservative bypass mechanism that exploits preexisting biases for homodinucleotides to increase the likelihood that the correct base is inserted opposite a non-instructive lesion. It will also be interesting to determine how HMCES-DPCs influence the mutation burden in non-small cell lung cancer and other cancers associated with elevated levels of AP sites^39^.

## Supporting information

Molecular dynamics simulation of AP site- or C3 spacer-containing DNA template in complex with DNA primer and REV3.

## Acknowledgements

We thank members of the Semlow lab for helpful discussions and comments on the manuscript. D.R.S. is supported by NIH grant no. R01GM151410 and a Shurl and Kay Curci Foundation Research Grant and is a Ronald and JoAnne Willens Scholar and a Vallee Scholar.

## Author contributions

X.H. performed the experiments described in Figure 1, Figure 2, C-F, Figure 3, B,C, and E, Figure 4, Supplementary Figure S1, A, C, and D, Supplementary Figure S2, A-D and G-L, and Supplementary Figure S3. X.H. and Y.C. performed the experiments described in Figure 5 and Supplementary Figure S4. K.T.N. performed the experiments described in Figure 3D and Supplementary Figure S2, E and F. T.C.X. and G.L. performed MD simulations described in Figure 6 and Supplementary Figure S5 under the supervision of W.A.G. D.R.S. performed the experiments described in Figure 2B and Supplementary Figure S1B. X.H., Y.C.X, W.A.G., and D.R.S. designed and interpreted the experiments. D.R.S. wrote the manuscript with input from all authors.

## Methods

Sequences of oligonucleotides mentioned in methods can be found in Supplementary Table 1. Next generation sequencing read counts can be found in Supplementary Tables 2–9. All unique materials are available from commercial suppliers or are available on request from the authors.

## Statistics and reproducibility

All experiments described were performed at least twice and representative results are shown.

### Preparation of *Xenopus* egg extracts

Animal work performed at Caltech was approved by the IACUC (Protocol IA20-1797 approved 28 May 2020). The institution has an approved Animal Welfare Assurance (no. D16-00266) from the NIH Office of Laboratory Animal Welfare. Preparation of high-speed supernatant (HSS) and nucleoplasmic extracts (NPE) from *Xenopus laevis* eggs was performed as described previously^40^. Briefly, HSS was prepared from eggs collected from six laboratory bred wild-type adult female *X. laevis* (aged >2 years). Eggs were dejellied in 1 L of 2.2% (w/v) cysteine, pH 7.7, washed with 2 L 0.5X Marc’s Modified Ringer’s solution (2.5 mM HEPES-KOH [pH 7.8], 50 mM NaCl, 1 mM KCl, 0.25 mM MgSO_4_, 1.25 mM CaCl_2_, 0.05 mM EDTA), and washed with 1 L of Egg Lysis Buffer (ELB) (10 mM HEPES-KOH [pH 7.7], 50 mM KCl, 2.5 mM MgCl_2_, 250 mM sucrose, 1 mM dithiothreitol (DTT), and 50 μg/mL cycloheximide). Eggs were then packed in 14-mL round-bottom Falcon tubes at 200 x g using a Sorvall ST8 swinging bucket rotor. Eggs were supplemented with 5 μg/mL aprotinin, 5 μg/mL leupeptin, and 2.5 μg/mL cytochalasin B and then crushed by centrifugation at 20,000 x g for 20 min at 4 °C in a TH13-6×50 rotor using a Sorvall Lynx 4000 centrifuge. The low-speed supernatant (LSS) was collected by removing the soluble extract layer and supplemented with 50 μg/mL cycloheximide, 1 mM DTT, 10 μg/mL aprotinin, 10 μg/mL leupeptin, and 5 μg/mL cytochalasin B. This extract was then spun in thin-walled ultracentrifuge tubes at 260,000 x g for 90 min at 2 °C in a TLS-55 rotor using a tabletop ultracentrifuge. Lipids were aspirated off, and HSS was harvested, aliquoted, snap frozen in liquid nitrogen, and stored at −80 °C. To prepare NPE, eggs were collected from 20 laboratory bred wild-type female *X. laevis* (aged >2 years). Eggs were dejellied in 2 L of 2.2% (w/v) cysteine, pH 7.7, washed with 4 L 0.5X Marc’s Modified Ringer’s solution (2.5 mM HEPES-KOH [pH 7.8], 50 mM NaCl, 1 mM KCl, 0.25 mM MgSO_4_, 1.25 mM CaCl_2_, 0.05 mM EDTA), and washed with 2 L of Egg Lysis Buffer (ELB) (10 mM HEPES-KOH [pH 7.7], 50 mM KCl, 2.5 mM MgCl_2_, 250 mM sucrose, 1 mM dithiothreitol (DTT), and 50 μg/mL cycloheximide). Eggs were then packed in 14-mL round-bottom Falcon tubes at 200 x g using a Sorvall ST8 swinging bucket rotor. Eggs were supplemented with 5 μg/mL aprotinin, 5 μg/mL leupeptin, and 2.5 μg/mL cytochalasin B and then crushed by centrifugation at 20,000 x g for 20 min at 4 °C in a TH13-6×50 rotor using a Sorvall Lynx 4000 centrifuge. The low-speed supernatant (LSS) was collected by removing the soluble extract layer and supplemented with 50 μg/mL cycloheximide, 1 mM DTT, 10 µg/mL aprotinin, 10 µg/mL leupeptin, 5 µg/mL cytochalasin B, and 3.3 μg/mL nocodazole. The LSS was then spun at 20,000 x g for 15 min at 4 °C in a TH13-6×50 rotor using a Sorvall Lynx 4000 centrifuge. The cytoplasm was transferred to a 50-mL conical tube. ATP regenerating mix (2 mM ATP, 20 mM phosphocreatine and 5 μg/mL creatine phosphokinase) was added to the extract. Nuclear assembly reactions were initiated by adding demembranated *X. laevis* sperm chromatin to a final concentration of 6,600 units/µL. After 75–90 min incubation, the nuclear assembly reactions were centrifuged for 3 min at 17,000 x g at 4 °C in a TH13-6×50 rotor using a Sorvall Lynx 4000 centrifuge. The nuclear layer was then harvested and spun at 260,000 x g for 30 min at 2 °C in a TLS-55 rotor using a tabletop ultracentrifuge. Finally, lipids were aspirated off, and NPE was harvested, aliquoted, snap frozen in liquid nitrogen, and stored at −80 °C.

### Protein expression and purification

Biotinylated LacI was expressed and purified essentially as described previously^19^. Briefly, LacI with a C-terminal AviTag and biotin ligase were coexpressed in T7 Express Cells supplemented with 50 µM biotin from pET11a[LacR-Avi] and pBirAcm (Avidity) vectors, respectively. AviTag-LacI and biotin ligase expression were induced with 1 mM isopropyl-β-d-thiogalactoside (IPTG) in media supplemented with 50 μM biotin to ensure efficient biotinylation of the AviTag-LacI. Cell pellets were lysed for 30 min on ice in lysis buffer containing 50 mM Tris-HCl (pH 7.5), 5 mM EDTA, 100 mM NaCl, 1 mM DTT, 10% sucrose (w/v), 1× cOmplete protease inhibitors, 0.2 mg/mL lysozyme, 0.1% Brij 58. The insoluble fraction was pelleted by centrifugation at 20,000g for 30 minutes at 4 °C in a A27-6×50 rotor using a Sorvall Lynx 4000 centrifuge. Chromatin-bound LacI was then suspended in 50 mM Tris-HCl (pH 7.5), 5 mM EDTA, 1 M NaCl, 30 mM IPTG, 1 mM DTT and released from the DNA by sonication followed by addition of polymin P to 0.03-0.06 % (w/v) at 4 °C. LacI was then precipitated with 37% ammonium sulfate, pelleted by centrifugation, and resuspended in buffer containing 50 mM Tris-HCl (pH 7.5), 1 mM EDTA, 2.6 M NaCl, 1 mM DTT and 1× cOmplete protease inhibitors. Next, biotinylated LacI was bound to with SoftLink avidin resin for 90 min at 4 °C, washed with 50 mM Tris-HCl (pH 7.5), 1 mM EDTA, 2.6 M NaCl, 1 mM DTT, 1× cOmplete protease inhibitors and eluted with 50 mM Tris-HCl (pH 7.5), 100 mM NaCl, 1 mM EDTA, 5 mM biotin and 1 mM DTT. Pooled fractions containing LacI were buffer exchanged into 50 mM Tris-HCl (pH 7.5), 150 mM NaCl, 1 mM EDTA, 1 mM DTT using an Amicon Ultra-0.5 mL 3K MWCO filter unit. LacI aliquots were snap frozen in liquid nitrogen and stored at −80 °C.

### Immunodepletions

Immunodepletions using antibodies raised against SPRTN, the N- or C-terminus of REV1, and Pol κ were performed as previously described^41–43^. Briefly, Protein A Sepharose Fast Flow (Cytiva) resin was washed with 1X PBS and incubated with 5 volumes of 1 mg/mL antibody overnight at 4 °C. The next day, the beads were washed twice with 1X PBS, once with ELB mix (2.5 mM MgCl_2_, 50 mM KCl, 10 mM HEPES, 250 mM sucrose), twice with ELB mix supplemented with 0.5 M NaCl, and 3 times with ELB mix. For SPRTN and Pol κ, depletions were performed by adding 5 volumes of egg extracts (HSS and NPE separately; 2 rounds for HSS and 3 rounds for NPE) to 1 volume of beads and incubating on a rotating wheel at room temperature for 20 min per round. For REV1, depletions were performed by adding 5 volumes of extract were added to 1 volume of beads and incubated at room temperature for 20 minutes per round. HSS was depleted with REV1-N beads for one round and REV1-C beads for one round, whereas NPE was depleted with REV1-N beads for two rounds and REV1-C beads for one round. Between rounds of depletion, the samples were spun at 622 x g for 30 seconds in an S-24-11-AT rotor in an Eppendorf 5430R centrifuge, and extract supernatants were collected.

### Immunoblotting

Samples in 1X Laemmli loading buffer (generally corresponding to 1-2 µL of replication reaction) were resolved on 10% or 4-15% acrylamide Mini-PROTEAN or Criterion TGX precast gels (Bio-Rad) and transferred to polyvinyl difluoride (PVDF) membranes (Thermo Fisher). Membranes were blocked with 5% nonfat milk in 1X PBS buffer with 0.05% (v/v) Tween 20 (PBST) for 1 hour at room temperature, rinsed 4 times with 1X PBST, and incubated with primary antibodies diluted to an appropriate concentration in 1X PBST at 4 °C overnight with shaking. Primary antibody dilutions were as follows: Rabbit polyclonal anti-SPRTN (*Xenopus*; Pocono Rabbit Farm and Laboratory, rabbit 31053), 1:5,000; Rabbit polyclonal anti-REV1 C terminus (*Xenopus*; Pocono Rabbit Farm and Laboratory rabbit 714), 1:5,000; Rabbit polyclonal anti-Pol κ (*Xenopus*; Biosynth, rabbit 7949), 1:5,000. The membranes were washed with 1X PBST 3 times for 10 minutes each at room temperature, then incubated for 30 minutes at room temperature with goat anti-rabbit peroxidase-conjugate secondary antibody (IgG, H + L; Jackson ImmunoResearch catalog no. 111-035-003) diluted 1:25,000 in 1X PBST with 5% (w/v) non-fat milk. After the incubation, the membranes were again washed with 1X PBST 3 times for 10 minutes each at room temperature, then incubated with ProSignal Pico Spray chemiluminescence substrate (Prometheus) for 1-2 minutes at room temperature and imaged using a ChemiDoc Imaging System (Bio-Rad). Goat anti-rabbit peroxidase-conjugate secondary antibody (IgG, H + L) was manufacturer validated by antigen-binding assay, western blotting and/or enzyme-linked immunosorbent assay (ELISA). All antibodies against *Xenopus* proteins used in this study were validated by western blotting using *Xenopus* egg extracts.

### Preparation of oligonucleotide duplexes with site-specific interstrand cross-links

Site-specific cross-links were prepared as previously described^16^. Oligonucleotides containing a site-specific deoxyuracil (dU) were annealed complementary oligonucleotides (5 µM each oligonucleotide) in 30 mM HEPES-KOH (pH 7.4), 100 mM NaCl by heating to 95 °C for 5 min, followed by cooling at 1 °C/min to 18 °C. The annealed duplex was ethanol-precipitated and resuspended in water at a final concentration of 5 µM. The duplex was then treated with 0.05 U/µL uracil-DNA glycosylase (NEB) in 1X UDG buffer (20 mM Tris-HCl, 10 mM DTT, 10 mM EDTA; pH 8.0) for 120 min at 37 °C, followed by extraction with phenol:chloroform:isoamyl alcohol (25:24:1; pH 8.0) and ethanol precipitation. The product was dissolved in 50 mM HEPES-KOH (pH 7.4), 100 mM NaCl at 5 µM and incubated at 37 °C for 7 days to 15 days allow cross-link formation. Cross-linked DNA duplexes were ethanol-precipitated, dissolved in 2x formamide loading buffer (86% formamide, 1X TBE, 20 mM EDTA) at 200 µM, and resolved on a 20% acrylamide/bis (19:1), 1x TBE, 8 M urea gel (1.5 mm thick). Gels were stained with SYBR Gold, and cross-linked products were visualized using a Blue-Light Transilluminator. The slower-migrating band corresponding to the cross-link was excised with a razor blade, crushed into fine pieces, and incubated with TE buffer (pH 8.0) at 4 °C for 18 h with rotation, with periodic replacement of the elution buffer. The eluates were combined, filtered through a 0.22 µm syringe-driven filter unit to remove gel debris, and concentrated by n-butanol extraction until the aqueous phase volume was <1 mL. A single round of chloroform extraction was then performed. The cross-linked duplexes were ethanol-precipitated, dissolved in 10 mM Tris-HCl (pH 8.5), flash-frozen in liquid nitrogen, and stored at –80 °C.

### Preparation of oligonucleotide duplexes with site-specific abasic sites

The uracil-containing oligonucleotide (5 µM) was treated with 0.05 U/µL uracil-DNA glycosylase (NEB) in 1× UDG buffer (20 mM Tris-HCl, 10 mM DTT, 10 mM EDTA; pH 8.0) for 120 min at 37 °C, followed by extraction with phenol:chloroform:isoamyl alcohol (25:24:1; pH 8.0) and ethanol precipitation. The resulting AP-oligonucleotide was resuspended in 10 mM Tris-HCl (pH 9.0). An equimolar amount of the complementary strand was then added, and the oligonucleotides were annealed at room temperature in 10 mM Tris-HCl (pH 9.0), 50 mM NaCl, and 0.1 mM EDTA at 5 µM for 15 min.

### Preparation of oligonucleotide duplexes with site-specific peptide adducts

The uracil-containing oligonucleotide (5 µM) was treated with 0.05 U/µL uracil-DNA glycosylase (NEB) in 1× UDG buffer (20 mM Tris-HCl, 10 mM DTT, 10 mM EDTA; pH 8.0) for 120 min at 37 °C, followed by extraction with phenol:chloroform:isoamyl alcohol (25:24:1; pH 8.0) and ethanol precipitation. The resulting AP-oligonucleotide was resuspended in 10 mM HEPES (pH 7.4), 5 mM NaCl, 5 mM DTT, 0.5 mM EDTA, and 0.1 to 0.5 mg/mL HMCES-derived peptides (GenScript) at a final concentration of 10 µM, and incubated at 37 °C for 2 h to allow cross-link formation. The reaction mixture was then extracted with phenol:chloroform:isoamyl alcohol (25:24:1; pH 8.0) and ethanol precipitated. The adduct oligonucleotide was resuspended in 10 mM Tris-HCl (pH 9.0), combined with an equimolar amount of complementary strand, and annealed at room temperature in 10 mM Tris-HCl (pH 9.0), 50 mM NaCl, and 0.1 mM EDTA at 5 µM for 15 min.

### Preparation of plasmids containing site-specific lesions

Plasmids containing engineered lesions were prepared as described previously^44^. Briefly, the backbone plasmid was digested with BbsI in NEBuffer r2.1 (10 mM Tris-HCl, 50 mM NaCl, 10 mM MgCl₂, 100 µg/mL recombinant albumin; pH 7.9) for 18 h at 37 °C, followed by extraction with phenol:chloroform:isoamyl alcohol (25:24:1; pH 8.0) and ethanol precipitation. The linearized plasmid was dissolved in TE (pH 8.0) and purified over a HiLoad 16/60 Superdex 200 column using isocratic flow of TE (pH 8.0). Fractions containing the digested plasmid were pooled, precipitated with isopropanol, and dissolved in 10 mM Tris-HCl (pH 8.5). Lesion-containing duplexes were ligated into the BbsI-digested backbone using 0.4 U/µL T4 DNA ligase (NEB) in 1× ligase buffer (50 mM Tris-HCl [pH 8.0], 10 mM MgCl₂, 1 mM ATP, 10 mM DTT) at 16 °C for 12h. The ligation products were concentrated using QIAGEN-tip 500 columns, and DNA was eluted with 1.25 M NaCl, 50 mM Tris-HCl (pH 9.0 for plasmids harboring a peptide adduct and pH 8.5 for other plasmids) in 1 mL fractions. Pooled fractions were buffer exchanged into TE (pH 9.0 for plasmids harboring a peptide adduct and pH 8.5 for other plasmids) with a 4 mL Amicon filter unit (3 kDa MWCO). CsCl was added to the DNA to a final solution density of 1.6 g/mL, and ethidium bromide was added to 50 µg/mL. The DNA was transferred to a Quick-Seal tube and centrifuged for 16 h at 4 °C in an NVT-90 rotor at 285,907 x g. The covalently closed circular plasmid was collected, extracted with water-saturated n-butanol to remove ethidium bromide, buffer-exchanged into TE (pH 9.0 for plasmids harboring a peptide adduct and pH 8.0 for other plasmids), and concentrated using a 4 mL Amicon filter unit (3 kDa MWCO). Aliquots were snap-frozen in liquid nitrogen and stored at –80 °C.

### Replication reactions

Plasmid replication reactions in *Xenopus* egg extracts were performed as previously described^45^. HSS was thawed and supplemented with nocodazole (3 ng/µL) and an ATP-regenerating system (ARS; 20 mM phosphocreatine, 2 mM ATP, and 5 ng/µL creatine phosphokinase), followed by centrifugation at 21,130 × g for 5 min at 4 °C in an Eppendorf 5424 R centrifuge. The clarified HSS was collected for plasmid licensing. Plasmids were added to a final concentration of 7.5 ng/µL and incubated at room temperature for 30 min to allow licensing. For assays involving LacR acting as a replication block, plasmids were pre-incubated with 14 µM biotinylated LacI for 1 hour at room temperature before being added to HSS. Where indicated, the licensing reaction was supplemented with 167–333 nM 3000 Ci/mmol [α-^32^P]dCTP. NPE was supplemented with 20 mM phosphocreatine, 2 mM ATP, 5 µg/mL creatine phosphokinase, and 4 mM DTT, then diluted 1:1 with egg lysis buffer (ELB) to generate a 50% NPE solution. Replication was initiated by mixing one volume of licensed plasmids with two volumes of 50% NPE. Where indicated, NPE mix was supplemented with 300 µM MG262.

### Base excision repair reactions

To examine base excision repair of AP sites and AP site analogs in egg extract, 10 µL aliquots of HSS licensing reactions were quenched with 100 µL of clear replication stop mix (50 mM Tris-HCl [pH 8.8], 0.5% SDS, 25 mM EDTA). Samples were digested with 200 µg/mL RNAse A for 30 mins at 37 °C and then with 1 mg/mL proteinase K for 60 mins at 37 °C. Samples were adjusted to 200 µL with 10 mM Tris-HCl (pH 8.8]), extracted twice with phenol:chloroform:isoamyl alcohol (25:24:1; pH 8.0), extracted once with chloroform, and ethanol precipitated. DNA was resuspended in 10 µL of 10 mM Tris-HCl (pH 8.5). 3 µL of recovered DNA was incubated with 1 U APE1 in 1X NEBuffer 4 (50 mM potassium acetate, 20 mM Tris-acetate, 10 mM magnesium acetate, 1 mM DTT; pH 7.9) in a total volume of 10 µL at 37 °C for 90 mins. Samples were resolved on a 0.8% agarose, 1X TBE, 0.5 µg/mL ethidium bromide gel and visualized with a ChemiDoc Imaging System (Bio-Rad).

### Native agarose gel analysis

For native agarose gel analysis, 1 µL aliquots of replication reaction were removed at the indicated time points and quenched in 6 µL replication stop mix (80 mM Tris-HCl [pH 8.0], 8 mM EDTA, 0.13% phosphoric acid, 10% Ficoll, 5% SDS, 0.1% bromophenol blue). After all samples were collected, proteinase K was added to 1.25 mg/mL, and reactions were incubated at 37 °C for 1 h. Replication intermediates and products were resolved on 0.8% agarose, 1x TBE gels, dried, and visualized by phosphorimaging. Autoradiograms were imaged with an Amersham™ Typhoon™ biomolecular imager (Cytiva) and analyzed using Image Lab v.6.4.0.

### Nascent strand analysis

Nascent strand analysis was performed essentially as previously described^46^. Replication reactions were initiated, and 4 µL aliquots were collected at the indicated time points and quenched in 40 µL clear replication stop mix (50 mM Tris-HCl [pH 8.8], 0.5% SDS, 25 mM EDTA). After all time points were collected, samples were digested sequentially with RNase A (200 µg/mL, 30 min, 37 °C) and then with proteinase K (1 mg/mL) at 37 °C for 1 h. Then the samples were adjusted to 200 µL with 10 mM Tris-HCl [pH 8.5], extracted twice with phenol:chloroform:isoamyl alcohol (25:24:1, pH 8.0) and once with chloroform, then ethanol-precipitated. DNA was resuspended in 10 µL of 10 mM Tris-HCl [pH 8.5] and stored at –20 °C. 2 µL of recovered DNA were incubated with a combination of 2 U AflIII and 4 U EcoRI or a combination of 5 U PstI and 5 U BamHI in 50 mM Tris-HCl (pH 8), 100 mM NaCl, 10 mM MgCl_2_ in a total volume of 5 µL at 37 °C for 4 hours. Reactions were stopped by adding 5-20 µL of Invitrogen Gel Loading Buffer II. Primers for sequencing ladders were radiolabeled by incubating 200 nM sequencing primer with 200 nM [γ-^32^P]ATP and 20 U T4 polynucleotide kinase (NEB) in 1X T4 PNK reaction buffer (70 mM Tris-HCl, 10 mM MgCl_2_, 5 mM DTT; pH 7.6) in a total volume of 50 µL for 30 minutes at 37 °C. T4 PNK was inactivated by heating at 95 °C for 2 minutes, and the unincorporated radionucleotides were removed using a Micro Bio-Spin column (Bio-Rad) per the manufacturer’s instructions. Sequencing ladders were made using the Thermo Sequenase Cycle Sequencing Kit: 0.5 pmol of radiolabeled primer were mixed with 175 ng of pCtrl, 2 µL of reaction buffer, and 2 µL of sequenase in a total volume of 17.5 µL. 4 µL of this reaction mix was then added to 4 µL of ddA, ddG, ddC, or ddT termination mixes, and the reactions were incubated on a PCR block at 95 °C for 3 minutes, followed by 50 cycles of 95 °C for 30 seconds, 60 °C for 60 seconds, and 72 °C for 90 seconds. 8 µL of Invitrogen Gel Loading Buffer II were added to each reaction, and the sequencing ladders were stored at −20 °C until use. Immediately before loading, all samples were heated at 75 °C for 5 minutes and snap-cooled in an ice water bath. DNA was resolved on a 7% acrylamide/bis (19:1), 8 M urea, 0.8X GTG buffer (71 mM Tris-HCl, 23 mM taurine, 0.4 mM EDTA) gel. Sequencing gels were dried and visualized by phosphorimaging as described above.

### Strand-specific Southern blotting

Strand-specific Southern blotting was performed as previously described^46^. Replication reactions were initiated, and 9 µL aliquots were collected at the indicated time points and quenched in 90 µL clear replication stop mix (50 mM Tris-HCl [pH 8.8], 0.5% SDS, 25 mM EDTA). After all time points were collected, samples were digested sequentially with RNase A (200 µg/mL, 30 min, 37 °C). Samples were split, with proteinase K (1 mg/mL) added to one, and incubated at 37 °C for 1h. Then the samples were adjusted to 200 µL with 10 mM Tris-HCl (pH 8.5), extracted twice with phenol:chloroform:isoamyl alcohol (25:24:1, pH 8.0) and once with chloroform, then ethanol-precipitated. DNA was resuspended in 10 µL of 10 mM Tris-HCl (pH 8.5) and stored at –20 °C. For restriction digestion, 3 µL of recovered DNA were incubated with 4 U AseI and 8 U XhoI in 1× NEBuffer r3.1 (100 mM NaCl, 50 mM Tris-HCl, 10 mM MgCl₂, 100 µg/mL recombinant albumin; pH 7.9) in a total volume of 5 µL at 37 °C for 4 h. Digested samples were mixed with an equal volume of Gel Loading Buffer II (Invitrogen) and stored at –20 °C. Immediately before loading, samples were heated at 75 °C for 5 min and snap-cooled in ice water. DNA was resolved on a 7%-10% acrylamide/bis (19:1), 8 M urea gel prepared in 0.8× GTG buffer (71 mM Tris-HCl, 23 mM taurine, 0.4 mM EDTA). Following electrophoresis, gels were transferred to filter paper, and DNA was transferred to Hybond-XL membrane (Cytiva) in 0.5× TBE at 0.4 A using a Trans-Blot SD semidry transfer cell (Bio-Rad). Membranes were washed with 4× SSC for 5 min and crosslinked with 120,000 µJ cm⁻² of 254 nm UV light. Prehybridization was carried out in 25 mL ULTRAhyb Ultrasensitive Hybridization Buffer (Invitrogen) for 12 h at 42 °C. The 105-mer top-strand probe was 5′-end radiolabeled by incubating 0.2 µM oligonucleotide with 0.2 µM [γ-^32^P]ATP (3,000 Ci/mmol) and 10 U T4 polynucleotide kinase in 1× NEB T4 PNK buffer for 30 min at 37 °C in a total volume of 50 µL. The reaction was heat-inactivated at 65 °C for 20 min and purified using a Micro Bio-Spin Column with Bio-Gel P-6 (Bio-Rad). The radiolabeled probe was added to the hybridization buffer and incubated with the membrane at 42 °C for 24 h. Membranes were washed twice with 2× SSC, 0.1% SDS at 42 °C for 5 min each, dried briefly on blotting paper, and visualized by phosphorimaging. Autoradiograms were imaged with an Amersham™ Typhoon™ biomolecular imager (Cytiva) and analyzed using Image Lab v.6.4.0.

### Preparation of next-generation sequencing libraries and NGS data processing

For next-generation sequencing (NGS), plasmids were replicated as described above. Following phenol/chloroform extraction and ethanol precipitation, samples were resuspended to approximately 2 ng/µL. 5 µL of recovered DNA were incubated with Nt.BstNBI in 1x NEBuffer r3.1 (100 mM NaCl, 50 mM Tris-HCl, 10 mM MgCl₂, 100 µg/mL recombinant albumin; pH 7.9) in a total volume of 50 µL at 55 °C for 1 h and then 85 °C for 20 minutes. 10 µL of each sample was run on a 0.8% agarose, 1X TBE, 0.5 µg/mL ethidium bromide gel and visualized to confirm replication intermediates and products are nicked. The remaining samples were combined with 500 nM each of forward and reverse primers (see Supplementary Table 1), 200 µM dNTPs, and 8 units of Phusion High-Fidelity DNA Polymerase (NEB) in 1× Phusion HF buffer to a total volume of 400 µL, which was distributed across four PCR tubes. Amplicons were generated by PCR with an initial denaturation at 98 °C for 30 seconds, followed by 18 cycles of 98 °C for 10 seconds, 55 °C for 30 seconds, and 72 °C for 30 seconds, and a final extension at 72 °C for 10 minutes. Amplified DNA was purified using the Zymo DNA Clean & Concentrator kit according to the manufacturer’s instructions and analyzed on a 0.8% agarose gel in 1x TBE, stained with SYBR Gold, to confirm amplification. Purified samples were submitted to Genewiz for next-generation sequencing (Amplicon-EZ, Illumina^®^ short-read NGS, 150–500 bp). Reads were processed using Jupyter Notebook.

### MD simulations

The cryo-EM structure of yeast Pol ζ holoenzyme complexed with duplex DNA (PDB 7S0T^25^) was used as the structural template for all simulations. The nascent DNA strand primer and template were mutated to correspond to the pICL^AP^ sequence context while the DNA backbone remained fixed (primer 5’-GGCAGAGAGA, template 3’-CCGTCTCTCTXCTC), where X denotes the template strand position that was replaced with an AP site or C3 spacer. Two systems were simulated, one AP site-containing and one C3 spacer-containing DNA-REV3 complex. The Schrödinger software Protein Preparation Workflow^47^ was used to check and protonate the AP site- and C3 spacer-modified DNA-REV3 complex structures. To improve simulation efficiency, REV7_A_, REV7_B_, POL31 and POL32_N_ subunits of the yeast Pol ζ cryo-EM structure (PDB 7S0T) were removed as they do not make direct contact with the duplex DNA^25^. The cryo-EM did not resolve the structure of 17 loop segments in REV3. Those less than 20 residues long were predicted and filled in using the Schrödinger Prime software^48^; those with longer missing loop segments, the REV7 binding region and the POL31 binding region, were not included in the simulation. 10 cycles of simulated annealing from 100 K to 600 K was performed to optimize the conformation of the added loops, while the remainder of the structure was fixed. Thus, the simulated REV3 model contains residues 20-400, and residues 661-1373. The DNA template strand position templating the incoming dNTP was replaced with an AP site or a C3 spacer. The free incoming dNTP was modified to a dATP, and the Ca^2+^ ion at the active site was substituted by a Mg^2+^ ion. The iron/sulfur cluster was removed from the complex. Three independent 500 ns simulations were performed for each system. All molecular dynamics and subsequent analysis were conducted at 310 K using GROMACS 2025.4 software^49^, and the figures were prepared using VMD^50^. The CHARMM-GUI^51^ was used to set up the simulation system, where the DNA-REV3 complex was solvated in a water box containing 0.05 M KCl with 12 Å padding, and neutralized to a net charge of 0. The parameterization procedure for noncanonical components, including the AP site, C3 spacer, and the free dATP, followed the protocol described by Usicik et al.^52^, with modifications to ensure compatibility with the CHARMM force field framework. This includes using CGenFF server^53^ for partial charge calculations and assigning CHARMM atom types. Mg^2+^ ion parameters were taken from Grotz et al^54^. The CHARMM36m force field^55^ was used for the protein and the DNA, while the TIP3P model^56^ was used for water. Electrostatic interactions were modelled using the PME method^57^, with the cut-off distance for non-bonded interactions set to 8 Å. Periodic boundary conditions were used for energy minimization and MD. The full system includes ∼ 220,000 atoms per periodic cell. In GROMACS 2025.4^49^, after an initial 2,000 steps energy minimization, the system was heated incrementally from 0 to 310 K over 500 ps using the NVT ensemble with harmonic restraints applied to the protein and DNA. Next, NPT MD was carried out using the Berendsen thermostat^58^ and C-rescale barostat^59^, while gradually releasing restraints over another 500 ps. All subsequent production MD simulations were conducted at 310 K with 1 atm pressure in the NPT ensemble using 2 fs time steps. The LINCS algorithm was applied to all covalent bonds involving hydrogen atoms.

---

**Supplementary Figure S1.**
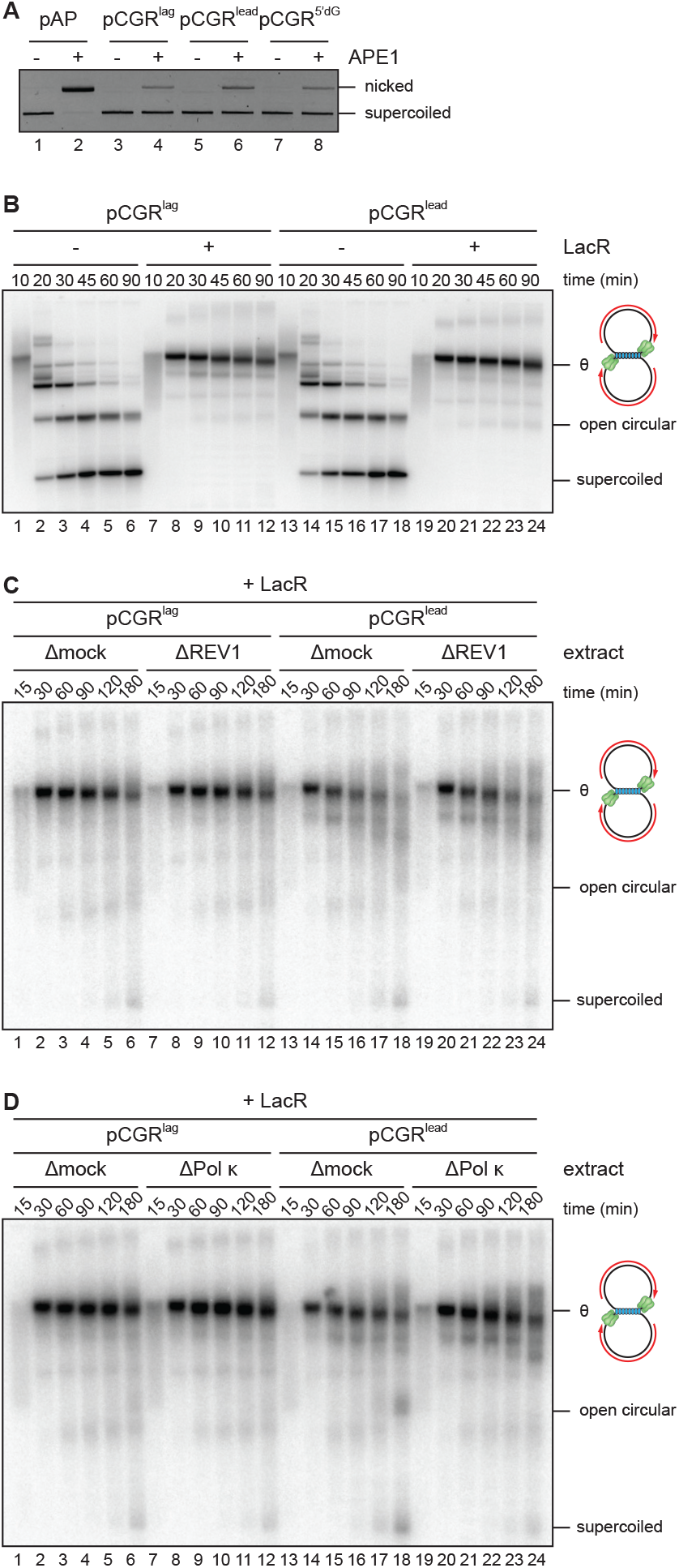
Replication of plasmids containing HMCES peptide adducts. **A**. CsCl gradient-purified plasmids containing AP sites or AP site-CGR tripeptide adducts were digested with recombinant APE1 and resolved on a native agarose gel containing ethidium bromide. Undigested plasmids resolve as a faster migrating supercoiled species while digested plasmids resolve as a slower migrating nicked species. Note that the adducted plasmids are largely resistant to nicking by APE1, indicating that most plasmids contain the adduct. **B-D**. The pCGR^lag^ and pCGR^lead^ replication reactions described in Figure 2B (**B**), Figure 2D (**C**), and Figure 2F (**D**) were deproteinized and replication intermediates and products were resolved on native agarose gels. Replication reactions were supplemented with [α-^32^P]dCTP, which is incorporated into the nascent DNA strands. Replication intermediates and products were visualized by autoradiography. Note that in the presence of LacR, replication results in the accumulation of θ intermediates that are produced when replication forks stall at the periphery of the 48x*lacO* array located on the plasmid.

**Supplementary Figure S2.**
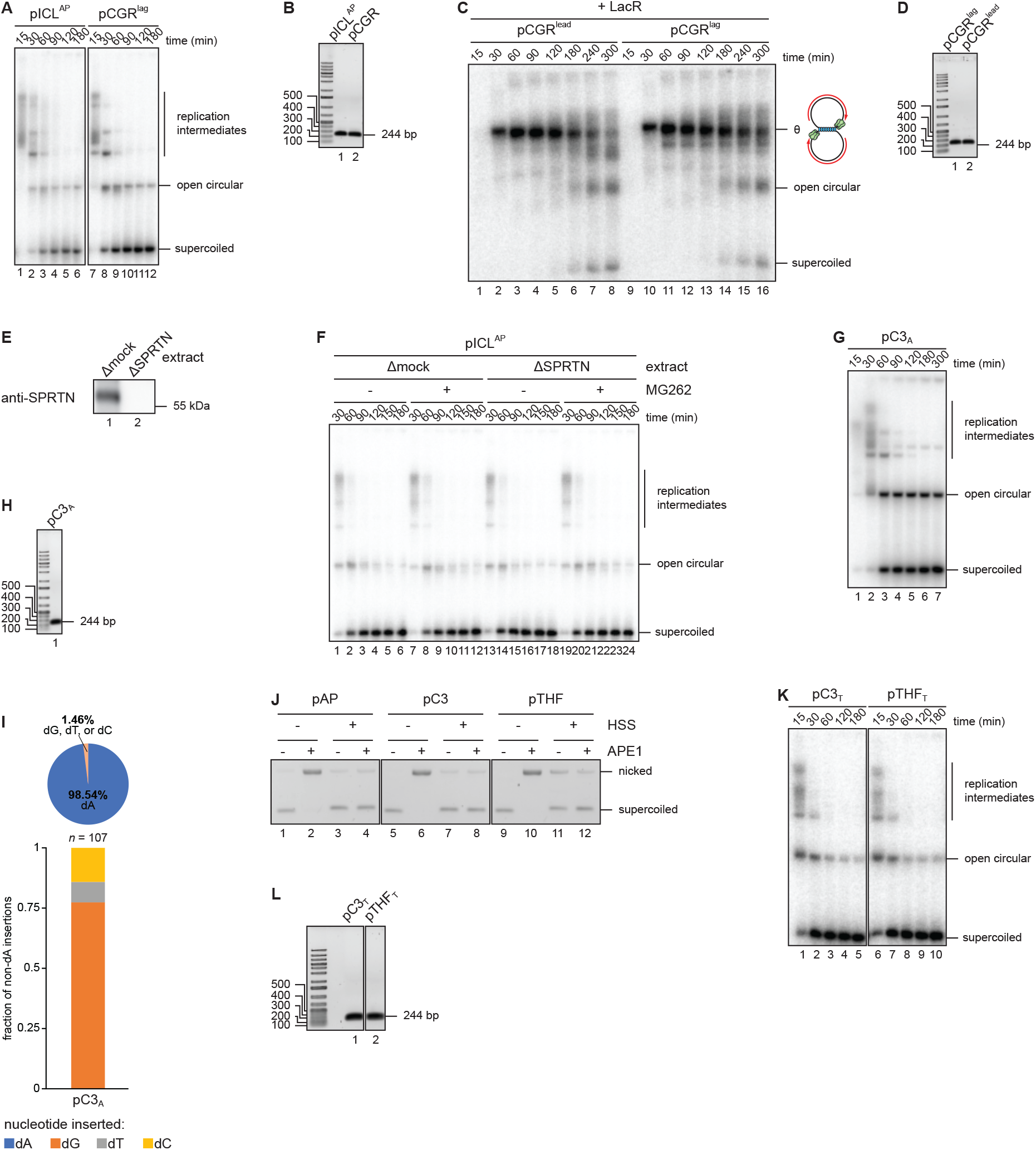
Analysis of HMCES adduct bypass products using next generation sequencing. **A**, **C**, **F**, **G**, and **K**. Aliquots of the replication reactions used to generate the sequencing libraries described in Figure 3B (**A**), Figure 3C (**C**), Figure 3D (**F**), **I** (**G**), Figure 3E (**K**) were supplemented with [α-^32^P]dCTP and replication intermediates and products were resolved on a native agarose gel and visualized as in Supplementary Figure S1B. **B**, **D**, **H**, and **L**. The PCR amplicon sequencing libraries described in Figure 3B (**B**), Figure 3C (**D**), **I** (**H**), and Figure 3E (**L**) were resolved on native agarose gels and visualized with SybrGold stain. Amplification of plasmid replication products produces a 244 bp fragment. **E.** SPRTN immunodepletion. The extracts used in the replication reactions described in Figure 3D and F were blotted for SPRTN. **I.** pC3_A_ was replicated in egg extract and replication products were sequenced as in Figure 3B. Top, pie chart depicting the fraction of reads corresponding to dA and non-dA insertions opposite the C3 spacer. Bottom, the fractions of non-dA reads corresponding to insertion of dG, dT, and dC opposite the C3 spacer is plotted. *n*, number of nascent strand non-dA insertion reads. **J**. pAP, pC3, and pTHF plasmids were incubated for 30 minutes in either buffer or high-speed supernatant (HSS) extract that is used to license DNA replication. The plasmids were recovered from extract and then digested with recombinant APE1. Both pAP and pC3 plasmids are efficiently nicked by APE1 following incubation in buffer, indicating that both the AP site and C3 spacer is cleaved by APE1. After incubation in HSS, both pAP and pC3 are refractory to cleavage by APE1, indicating that both the AP site and C3 spacer is efficiently removed in extract.

**Supplementary Figure S3.**
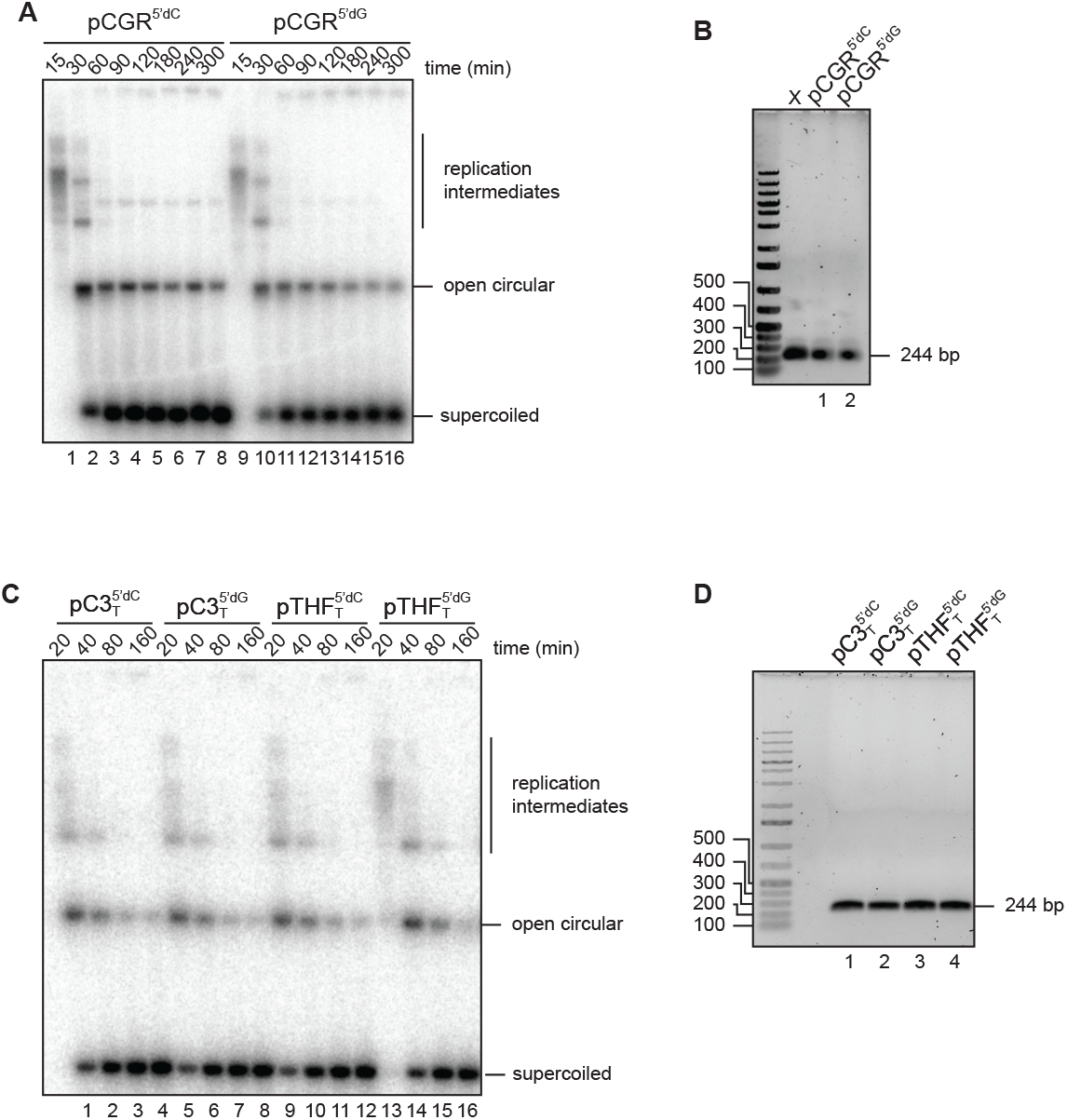
Analysis of 5’ template nucleotide effects during HMCES adduct bypass by next generation sequencing. **A** and **C**. Aliquots of the replication reactions used to generate the sequencing libraries described in Figure 4B (**A**) and Figure 4C (**C**) were supplemented with [α-^32^P]dCTP and replication intermediates and products were resolved on a native agarose gel and visualized as in Supplementary Figure S1B. **B**, and **D**. The PCR amplicon sequencing libraries described in Figure 4B (**B**) and Figure 4C (**D**), were resolved on a native agarose gel and visualized with SybrGold stain. Amplification of plasmid replication products produces a 244 bp fragment.

**Supplementary Figure S4.**
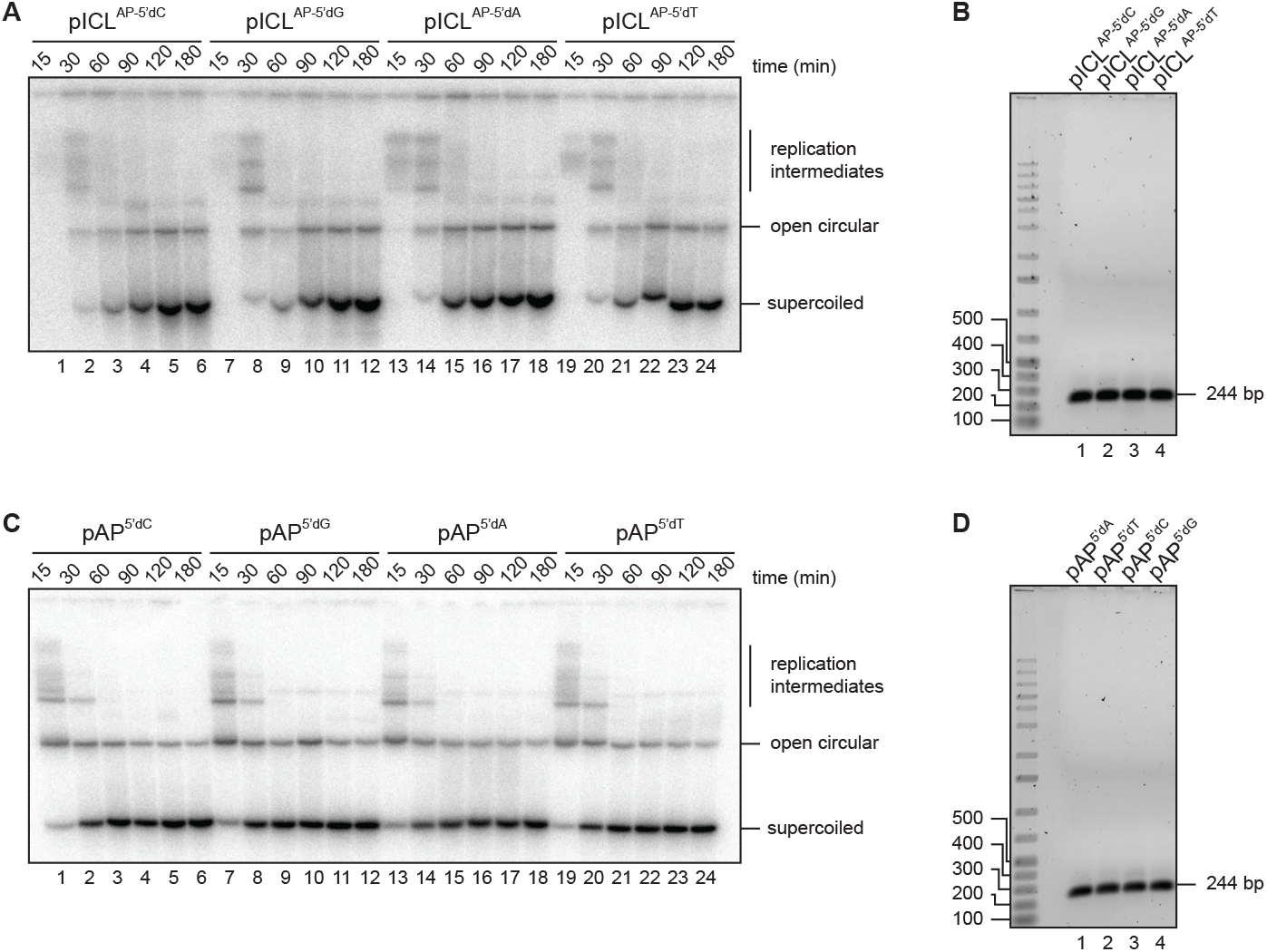
Analysis of 5’ template nucleotide effects during endogenous HMCE-DPC bypass by next generation sequencing. **A** and **C**. Aliquots of the replication reactions used to generate the sequencing libraries described in Figure 5A (**A**) and Figure 5B (**C**) were supplemented with [α-^32^P]dCTP and replication intermediates and products were resolved on a native agarose gel and visualized as in Supplementary Figure S1B. **B**, and **D**. The PCR amplicon sequencing libraries described in Figure 5A (**B**) and Figure 5B (**D**), were resolved on a native agarose gel and visualized with SybrGold stain. Amplification of plasmid replication products produces a 244 bp fragment.

**Supplementary Figure S5.**
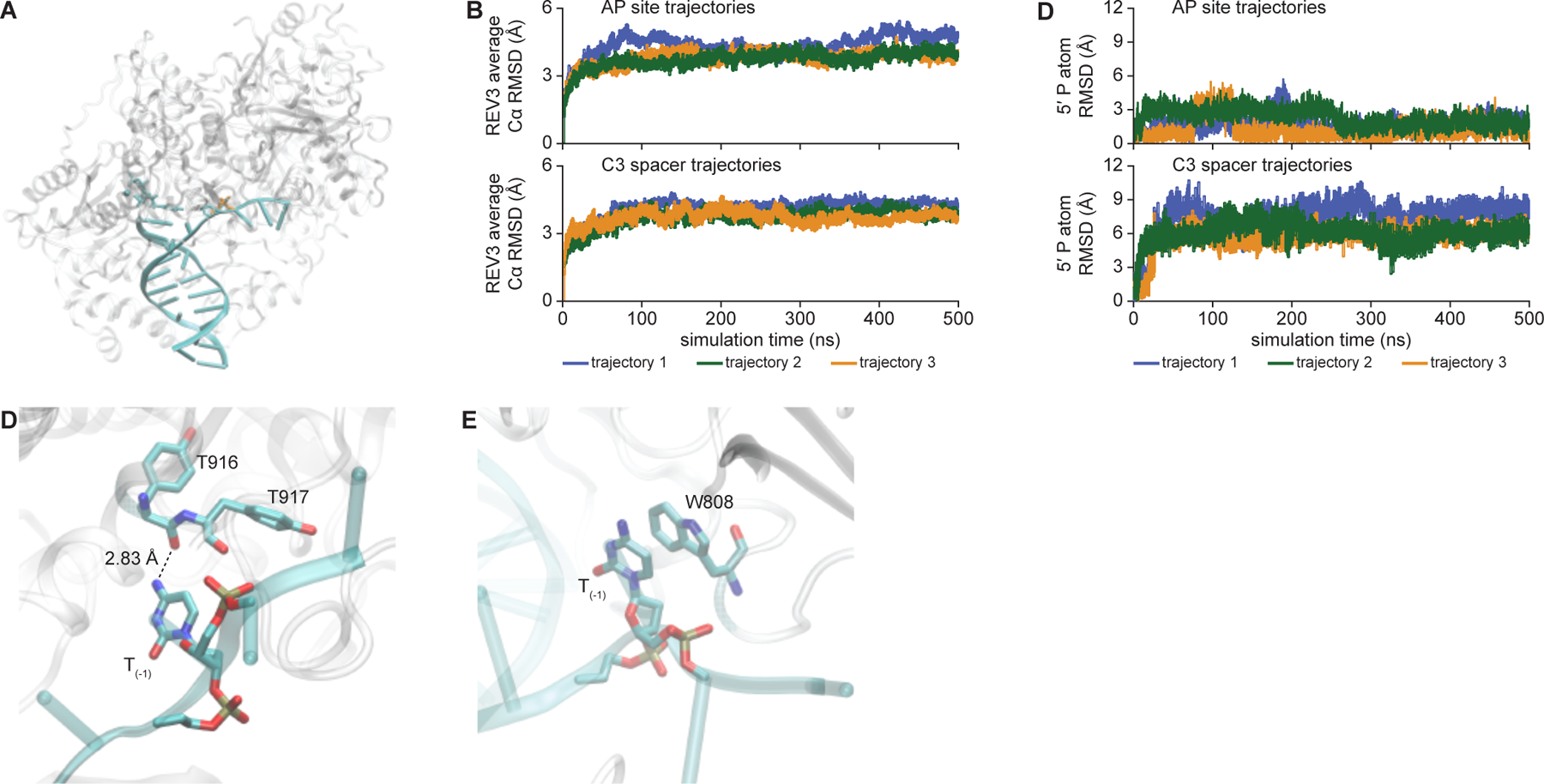
Molecular dynamics simulations of REV3-DNA complexes. **A.** Overlay of initial REV3-DNA structures for AP site and C3 spacer templates. Gray, yeast REV3 subunit; cyan, AP site DNA; gold, C3 spacer DNA. **B.** Trajectories for the average REV3 backbone Cα atom RMSD relative to the starting structure. **C.** Trajectories for the 5’ phosphorous atom RMSD relative to the starting structure. **D** and **E**. Rotation of C3 spacer 5’ template dC nucleotide (T_-1_) is stabilized by interactions with REV3. dC hydrogen bonding with T916 (**D**) and dC π-π stacking with W808 (**E**) are shown. Gray, yeast REV3 subunit; cyan, DNA duplex.

